# An early mechanosensitive window in bone fracture healing shapes long-term repair

**DOI:** 10.64898/2026.09.25.754061

**Authors:** Liesbeth Ory, Gabriele Nasello, Edoardo Borgiani, Tom Verbraeken, Kathleen Bosmans, Carla Geeroms, Inge Van Hoven, Elena Nefyodova, Claire Chabot, Liesbet Geris, Przemko Tylzanowski

**Author notes:** Corresponding author: Liesbet Geris. Liesbet Geris and Przemko Tylzanowski share senior authorship. Contributions: LO, PT, and LG conceptualized and designed the study. LO, GN, EB, and TV developed and adapted code. LO performed the experiments and analyzed the data. GN provided the Docker working environment. KB assisted with surgery, CG performed micro-CT acquisition, IVH assisted with single-cell isolation, and EN performed histological sectioning and staining. CC performed mechanical testing. LO wrote the first draft of the manuscript, and all other authors contributed to manuscript revision. All authors approved the submitted version.

## Abstract

Mechanical stability critically influences bone fracture healing, yet how the early mechanical environment directs the transition from inflammation to regeneration remains unclear. Using a murine femoral osteotomy model, we compared rigid, semirigid, and dynamically adjusted semirigid-to-rigid fixation. Semirigid fixation delayed healing on day 21, whereas a strategy of early compliance followed by increased stiffness after 7 days restored bridging and improved bone regeneration beyond constant rigid fixation, pointing to a critical early mechanosensitive window after fracture. Single-cell RNA sequencing across the first week, comparing semirigid with rigid fixation, showed that fixation stiffness altered intercellular signaling within one day of injury. Signaling among myeloid populations was more broadly increased under semirigid fixation, with monocytes the single exception, and signaling associated with resolution of inflammation was reduced. From day 5, periosteal and skeletal progenitor populations increasingly directed cartilage-associated matrix and growth factor signaling toward macrophages and chondrocytes, accompanied by increased *Sox9* regulon activity and increased COL10+ hypertrophic cartilage matrix by day 7. Reduced fixation stiffness therefore promotes a chondrogenic trajectory of repair, but increasing stiffness is required to redirect this response toward bone formation. *Piezo1* was expressed across the responding populations and shifted between compartments as healing progressed. PIEZO1 activation with Yoda1 under rigid fixation enhanced bone formation but provided no additional benefit under semirigid or dynamically adjusted fixation, whereas inhibition of mechanosensitive signaling with GsMTx4 impaired repair across all conditions. Together, these findings establish the early mechanical environment as a determinant of fracture-healing trajectory, shaping the transition from inflammation to regeneration, and show that successful repair depends not on maximal stability but on when stiffness is applied. Increasing either early mechanical stimulation or cellular mechanosensitivity enhanced regeneration, supporting mechanosensitive signaling as a key component of the early healing response.

## Introduction

Fracture healing requires a degree of mechanical stimulation, yet too much movement prevents it. Clinically, optimizing fixation strategies is therefore essential, as insufficient or excessive stability disrupts the balance between mechanical stimulation and biological progression, leading to delayed healing or non-union^1^. Bone fracture healing proceeds through sequential phases of inflammation, repair, and remodeling. Immediately after fracture, disruption of blood vessels results in hematoma formation at the injury site, creating a provisional microenvironment that initiates the inflammatory phase of healing. This fracture hematoma serves as an early signaling niche in which immune cells, including neutrophils, monocytes, and macrophages, are recruited and activated. These cells contribute to debris clearance as well as release cytokines and growth factors regulating the recruitment, proliferation, and differentiation of skeletal stem and progenitor cells. The progenitor cells subsequently differentiate toward chondrogenic and osteogenic lineages, forming a cartilage-rich callus that serves as a template for subsequent bone formation through endochondral ossification. Throughout these processes, mechanical cues play a pivotal but underappreciated role in regulating cellular behavior, matrix deposition, and vascularization^2^. Understanding how the early mechanical environment affects fracture healing is therefore of both biological and translational importance^1^.

The mechanical environment at the fracture site is largely determined by fixation strategy and evolves throughout the healing process. Fixation influences the interfragmentary movement occurring during loading, which determines the local tissue deformation and stresses experienced within the fracture gap. These mechanical cues influence cellular processes including migration, proliferation, differentiation, and extracellular matrix production, contributing to tissue formation and organization^3^. In particular, the local combination of tissue strain and hydrostatic pressure influences the differentiation of multipotent progenitor cells towards different tissue types, including fibrous tissue, fibrocartilage, or hyaline cartilage^4^. As the newly formed tissue progressively increases in stiffness, the mechanical environment changes accordingly, reducing interfragmentary deformation and creating conditions that favor ossification and bone formation^5^. A key unresolved question is therefore how the mechanical environment is interpreted by cells to determine regenerative trajectories. Interestingly, the biological response to mechanical stimulation depends not only on the magnitude of loading but also on its timing during repair. Early excessive instability disrupts tissue organization, whereas insufficient mechanical stimulation may delay bone formation. Indeed, dynamically increasing fixation stiffness during the first week after fracture has been shown to rescue delayed healing, pointing to a critical mechanosensitive window early in repair^6^. These findings suggest that the early mechanical environment influences healing not only through tissue stabilization but also by regulating cellular processes that determine regenerative progression. Recent spatially resolved approaches have begun to relate transcriptional responses directly to the local mechanical environment within a fracture site by combining micro-computed tomography, spatial transcriptomics and finite element analysis, demonstrating that cells in regions of high and low strain differ transcriptionally^7^. How the mechanical environment established by fixation shapes these responses over the course of early repair, however, remains unresolved.

Although biomechanical regulation and inflammation have traditionally been studied as separate aspects of fracture healing, growing evidence indicates that they are closely interconnected. The early inflammatory phase has emerged as a decisive regulator of regenerative outcomes. Immune cells recruited after injury do not merely clear damaged tissue but actively shape subsequent repair through cytokine production, regulation of angiogenesis, and interactions with skeletal progenitors^8^. Immune cells are also highly responsive to the mechanical properties of their environment, so fixation-dependent differences in interfragmentary movement may influence inflammatory trajectories directly^9^. What remains unclear is how these effects evolve within the phase where they begin. In murine fracture models the inflammatory response resolves over the first week, soft callus formation peaks around day 14, and mineralization and remodeling follows over the subsequent weeks. The first week therefore contains the transition from inflammation to early anabolic activity, during which the populations present, the signals they exchange, and the matrix they encounter change substantially^1,5^. Whether the mechanical environment acts at a single point in this transition or reshapes it progressively has not been established, and distinguishing these possibilities requires following intercellular signaling across the inflammatory phase rather than at a single timepoint. While prior work has established that the timing of mechanical stimulation shapes healing outcome at the tissue level, the cellular and molecular events unfolding within this mechanosensitive window, as well as any pathway linking fixation stiffness to those events, have not been resolved.

Mechanotransduction, the process by which cells convert mechanical stimuli into biochemical signals, provides a molecular link between the mechanical environment and cellular responses. Among the mechanosensitive pathways identified to date, PIEZO1 has emerged as a key mechanosensor in skeletal and immune biology, integrating mechanical cues with downstream processes including inflammatory signaling, osteogenic differentiation, and extracellular matrix deposition^10,11^. Beyond its ability to detect mechanical stimuli, PIEZO1 regulates cellular adaptation through calcium influx and downstream signaling cascades that influence inflammatory responses, tissue remodeling, and cell fate decisions. Through these mechanisms, PIEZO1 can modulate responses to mechanical stimuli across multiple cell types^12,13^. However, whether PIEZO1 functions as a molecular interpreter of fixation-dependent mechanical cues during the early inflammatory phase of fracture healing remains unknown.

Here we investigated how the early mechanical environment shapes fracture repair. We compared rigid, semirigid and dynamically adjusted semirigid-to-rigid fixation, using single-cell RNA sequencing across the first week to characterize how intercellular signaling responds to fixation stiffness. We then tested whether PIEZO1, which is expressed across the responding populations, contributes to these effects by pharmacologically modulating its activity under each fixation condition.

## Results

### Delayed stiffening improves healing and alters early callus composition

To investigate how the mechanical environment influences fracture repair, we applied rigid, semirigid, or dynamically adjusted semirigid-to-rigid fixators to murine femoral defects (Figure 1a). Micro-computed tomography (micro-CT) and histological analyses at 21 days post-fracture revealed distinct healing trajectories. Rigid fixation resulted in bone bridging, with a median bridged bone percentage of 39.1% and limited residual cartilage at the defect site (9.8 ± 2.1%). Semirigid fixation resulted in delayed healing, characterized by reduced bone bridging (median 26.8%, p=0.004) and increased cartilage persistence (21.6 ± 4.8%), as quantified by Alcian Blue staining (Figure 1b–c). In contrast, increasing fixation stiffness after 7 days resulted in enhanced bone bridging (median 53.5%, p<0.001 versus semirigid, p=0.004 versus rigid) compared with both semirigid and rigid fixation, and the lowest residual cartilage of the three conditions (4.7 ± 1.5%; Welch ANOVA p = 0.011 across the three groups), although pairwise comparisons after Holm correction did not individually reach significance (p = 0.055–0.065) (Figure 1c).

**Figure 1.**
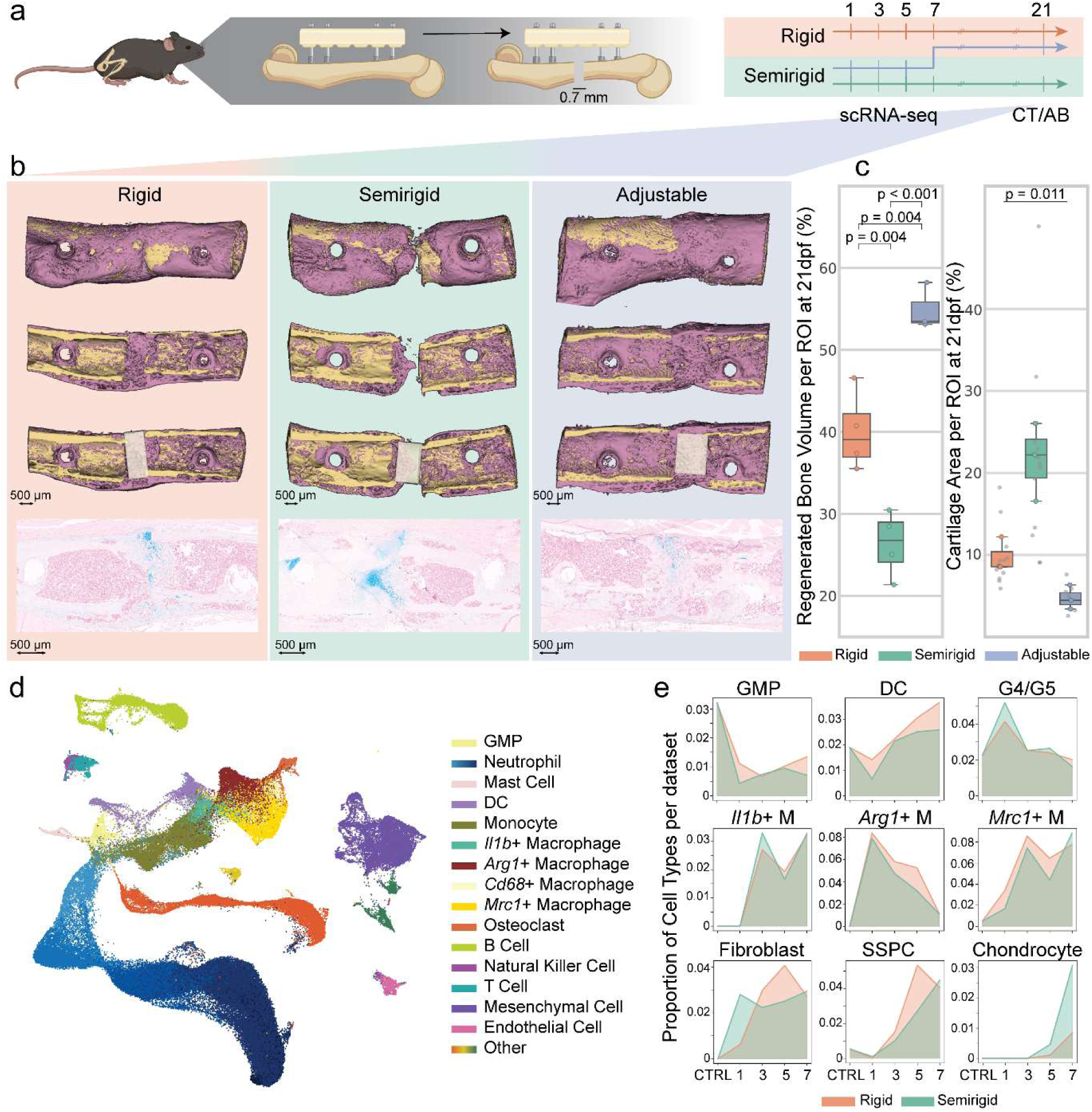
Delayed stiffening improves healing and alters cellular dynamics in the mouse osteotomy model. **a**, Schematic of the mouse long bone osteotomy model stabilized by three fixation strategies: rigid fixation, semirigid fixation, and the dynamically adjusted semirigid-to-rigid strategy. Samples were collected on days 1, 3, 5 and 7 for scRNA-seq, and on day 21 for CT analysis and Alcian Blue staining. **b**, Day 21 radiological and histological outcomes. Representative CT images (cortical bone in yellow, regenerated bone in purple) and corresponding Alcian Blue-stained histological sections (bottom row) of calluses under rigid, semirigid and dynamically adjusted fixation. The white box demarcates the ROI. **c**, Quantitative healing outcomes on day 21. Left, regenerated bone volume per ROI; boxes show median and interquartile range, points show individual animals (n = 4 rigid, 4 semirigid, 3 dynamically adjusted); one-way ANOVA with Tukey’s honestly significant difference test. Right, cartilage area per ROI; boxes show median and interquartile range of per-animal means, large points show per-animal means and small gray points individual sections (three to five per animal); n = 3 animals per group. Groups were compared by Welch ANOVA (p=0.011, shown on the panel); pairwise Welch’s t-tests with Holm correction, p = 0.055 (semirigid versus stiffening), 0.065 (rigid versus semirigid), 0.065 (rigid versus stiffening). **d**, UMAP of the pooled scRNA-seq dataset from all timepoints and fixation groups, colored by major cell type. **e**, Area plots showing proportional changes of specific cell types across control samples and days 1, 3, 5 and 7, comparing rigid (orange) and semirigid (green) fixation. scRNA-seq: single-cell RNA sequencing, CT: computed tomography, AB: Alcian Blue, ROI: region of interest, GMP: Granulocyte-Monocyte Precursor, DC: Dendritic Cell, SSPC: Skeletal Stem and Progenitor Cell.

Mechanical testing of the fixator constructs showed stiffnesses of 6.51 and 20.38 N/mm for the semirigid and rigid systems, respectively, and 4.33 and 11.41 N/mm for the dynamically adjusted system before and after stiffening (Supplementary Table 1). The semirigid system is thus more mechanically compliant, deforming more under a given load. The dynamically adjusted construct was the most compliant of all during the first week and, even after stiffening, remained well below the rigid system, yet this group achieved the greatest bone bridging. Because the adjustable and non-adjustable systems differ in absolute stiffness, this comparison evaluates a strategy of early compliance followed by increased stability, rather than isolating the timing of stabilization as a single variable.

To investigate the cellular mechanisms, we performed scRNA-seq on cells isolated from the early fracture-healing environment at days 1, 3, 5, and 7. Samples were collected from defects stabilized with either rigid or semirigid fixation, with contralateral intact bone collected as a control. This sampling captures cells within the hematoma and developing soft callus, together with cells present in the adjacent bone marrow, providing a broad representation of the cellular populations involved in early bone regeneration. UMAP visualization of the integrated dataset identified expected clusters corresponding to major cell populations involved in fracture healing. These included inflammatory populations dominated by myeloid cells, including myeloid progenitors, neutrophils, monocytes, and macrophages, as well as smaller lymphoid populations consisting of B and T cells. Mesenchymal and stromal populations involved in tissue regeneration were also identified (Figure 1d, Supplementary Figure 1).

Temporal analysis revealed dynamic changes in cellular composition during the first 7 days of healing. By day 1 after fracture, activated neutrophil and inflammatory macrophage populations, marked by *Arg1*, were already prominent. This was followed by a transition toward reparative macrophage states marked by *Mrc1* expression^14^.

Mesenchymal populations, including fibroblasts, were present from day 1, whereas skeletal stem and progenitor cell (SSPC) populations emerged at later time points. By day 7, increased abundance of chondroprogenitor and chondrocyte populations indicated progression toward cartilage formation (Figure 1e).

Fixation stiffness therefore shaped both the healing outcome on day 21 and the cellular composition of the callus during the first week.

### Semirigid fixation redirects early inflammatory signaling

Next, to dissect how the early mechanical environment influences immune responses, we examined the immune populations in this dataset in more detail. UMAP visualization of combined timepoints, using more granular cell-type annotations, identified distinct immune populations, including monocytes, neutrophils at different maturation states, macrophages, dendritic cell populations (including plasmacytoid, monocyte-derived, Clec9a+, and migratory dendritic cells), lymphoid populations including B cells at different activation states, *CD4*+ and *CD8*+ T cells, natural killer cells, and fibrocytes (Figure 2a, Supplementary Figure 1). Macrophage populations were further resolved into distinct transcriptional states, including *Il1b*-expressing, *Arg1*-expressing and *Mrc1*-expressing macrophage populations.

**Figure 2.**
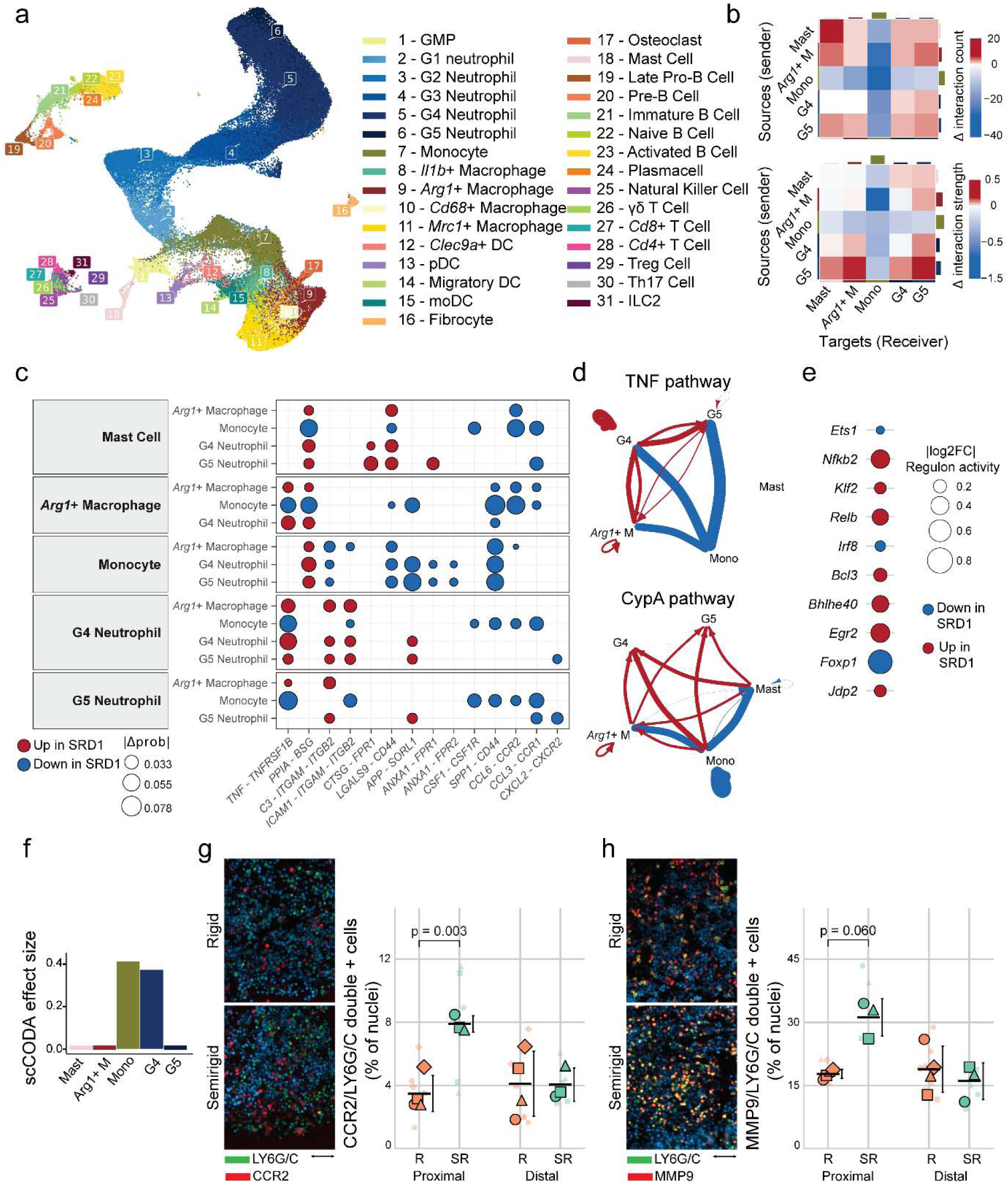
Fixation stiffness alters immune cell regulatory activity and intercellular communication during early fracture healing. **a**, UMAP of the integrated single-cell RNA-sequencing dataset showing immune cell populations identified across the fracture callus, colored by annotated cell type and state. **b**, Heatmaps showing the differential number (top) and differential strength (bottom) of predicted cell–cell interactions between sender and receiver populations at day 1 under semirigid compared with rigid fixation. Red indicates an increase and blue a decrease under semirigid fixation. **c**, Differential ligand–receptor interactions between the myeloid populations most active during the inflammatory phase at day 1. Senders are indicated at left, receivers on the vertical axis. Dot color indicates interactions increased (red) or decreased (blue) under semirigid fixation; dot size indicates the absolute difference in communication probability. **d**, Differential communication networks for the TNF (top) and CypA (bottom) pathways between the same populations. Arrows run from sender to receiver, width is proportional to the difference in communication probability, and color indicates direction as in **c**. **e**, Differential regulon activity between semirigid and rigid fixation at day 1. Dot size indicates the absolute log2 fold change; color indicates increased activity under semirigid (red) or rigid (blue) fixation. **f**, Compositional analysis (scCODA) at day 1, showing the myeloid populations displayed in **c**. The model was fitted across all annotated cell populations; effects were considered credible at a false discovery rate of 0.2, with Th17 Cell as reference. **g**, Immunofluorescence of the fracture callus at day 1 under rigid and semirigid fixation, stained for LY6G/C with CCR2. CCR2+LY6G/C+ cells as percentage of nuclei in proximal and distal regions under rigid (R) and semirigid (SR) fixation. Small symbols show individual regions of interest (two sections per animal) and large symbols animal means, with symbol shape identifying the animal; horizontal lines show group means with SD. Welch’s t-test per region, Holm-corrected for two comparisons. n = 4 rigid and 3 semirigid animals. Scale bar, 50 µm. **h,** Immunofluorescence of the fracture callus at day 1 under rigid and semirigid fixation, stained for LY6G/C with MMP9. MMP9+LY6G/C+ cells as percentage of nuclei in proximal and distal regions under rigid (R) and semirigid (SR) fixation. Small symbols show individual regions of interest (two sections per animal) and large symbols animal means, with symbol shape identifying the animal; horizontal lines show group means with SD. Welch’s t-test per region, Holm-corrected for two comparisons. n = 4 rigid and 3 semirigid animals. Scale bar, 50 µm. n = 4 to 5 biological replicates per condition and timepoint for all single-cell analyses. SRD1: semirigid day 1, RD1: rigid day 1, GMP: Granulocyte-Monocyte Progenitor, DC: Dendritic Cell, ILC2: Innate Lymphoid Cell 2, Mono: Monocyte, Mast: Mast Cell, Osteo-CAR: Osteoblast-CXCL12 Abundant Reticular Cell.

Gene regulatory network analysis of immune cells at day 1 revealed increased activity of regulons associated with inflammatory signaling under semirigid fixation, including the NF-κB-related regulators *Nfkb2*, *Relb* and *Bcl3*, together with *Bhlhe40* and *Jdp2* (Figure 2e). By day 3 these regulons showed reduced activity, with few differences remaining by day 7 (Supplementary Figure 2).

Cell–cell communication was inferred from ligand and receptor co-expression using CellChat, which returns a probability that an interaction is active rather than a direct measurement. Differences between conditions are therefore reported as differences in predicted communication. Comparison of cell–cell communication at day 1 among the myeloid populations most active during the inflammatory phase, mast cells, *Arg1*-expressing macrophages, monocytes, and G4 and G5 neutrophils, showed that both the number and strength of interactions were increased under semirigid fixation across most populations. Monocytes were the single exception, showing reduced interaction number and strength as both sender and receiver (Figure 2b).

Ligand–receptor analysis showed that this pattern was directional rather than uniform (Figure 2c). Interactions among mast cells, neutrophils and *Arg1*-expressing macrophages were increased under semirigid fixation, including TNF–TNFRSF1B, PPIA– BSG, C3–ITGAM–ITGB2, ICAM1–ITGAM–ITGB2 and LGALS9–CD44. In contrast, signaling directed toward monocytes was consistently reduced, including CCL6–CCR2, CCL3– CCR1 and CXCL2–CXCR2 chemokine signaling together with CSF1–CSF1R, SPP1–CD44 and APP–SORL1. Two features of this pattern emerged. TNF signaling was increased through TNFRSF1B between *Arg1*-expressing macrophages and neutrophils but reduced toward monocytes, indicating a shift in which cells receive TNF rather than an overall change in TNF availability (Figure 2c–d). Signaling through the formyl peptide receptors changed in ligand rather than in magnitude: ANXA1–FPR1 and ANXA1–FPR2 signaling from monocytes toward neutrophils was reduced, while CTSG–FPR1 signaling from mast cells toward the same populations was increased.

Compositional analysis showed increased proportions of monocytes and G4 neutrophils under semirigid fixation at day 1, with no credible change in the other populations examined (Figure 2f). Monocytes were therefore more abundant while being the only population with reduced communication, suggesting that the difference between fixation conditions lies in how myeloid cells are engaged rather than in how many are present.

To validate these single-cell findings, we performed immunofluorescence staining at day 1 for LY6G/C together with CCR2, marking inflammatory monocytes, and together with MMP9, marking activated neutrophils (Figure 2g,h). Consistent with the increased monocyte proportion in the compositional analysis, CCR2 and LY6G/C double positive cells were more than twice as frequent under semirigid fixation in the proximal region of the fracture (7.9 ± 0.5% versus 3.5 ± 1.1% of nuclei, p = 0.003), whereas no difference was observed distally (Figure 2g). The same proximal pattern was seen for LY6G/C+ cells overall (p = 0.046) and, less pronounced, for CCR2+ cells (p = 0.083) (Supplementary Figure 3a,b). MMP9 and LY6G/C double-positive cells showed the same directional difference proximally, without reaching significance (p = 0.060) (Figure 2h, Supplementary Figure 3a,b).

Thus, semirigid fixation reshaped the early inflammatory environment through coordinated changes in myeloid cell abundance and intercellular signaling, providing a distinct cellular context for subsequent tissue regeneration.

### Progenitor-derived signaling shifts toward chondrogenic support by day 7

Next, we asked whether these differences in the early inflammatory environment were accompanied by changes in skeletal progenitor populations and their signaling interactions during tissue formation. UMAP visualization identified distinct mesenchymal populations, including periosteal SSPCs (pSSPCs), bone marrow SSPCs (bmSSPCs), multiple fibroblast subtypes and osteochondral populations (Figure 3a). Across the four timepoints, rigid fixation showed a higher proportion of cells predicted to be multipotent (58.9% versus 47.2%), while chondrocyte-associated populations were more represented under semirigid fixation, including resting chondrocytes (0.3% to 2.0%), hypertrophic chondrocytes (7.2% to 14.6%) and osteoblast-committed chondrocytes (2.8% to 5.7%) (Figure 3b). These differences were most apparent on days 5 and 7 (Supplementary Figure 4). Given the limited abundance of several differentiated populations, these observations are descriptive trends in cellular distribution rather than statistically supported shifts in lineage commitment.

**Figure 3.**
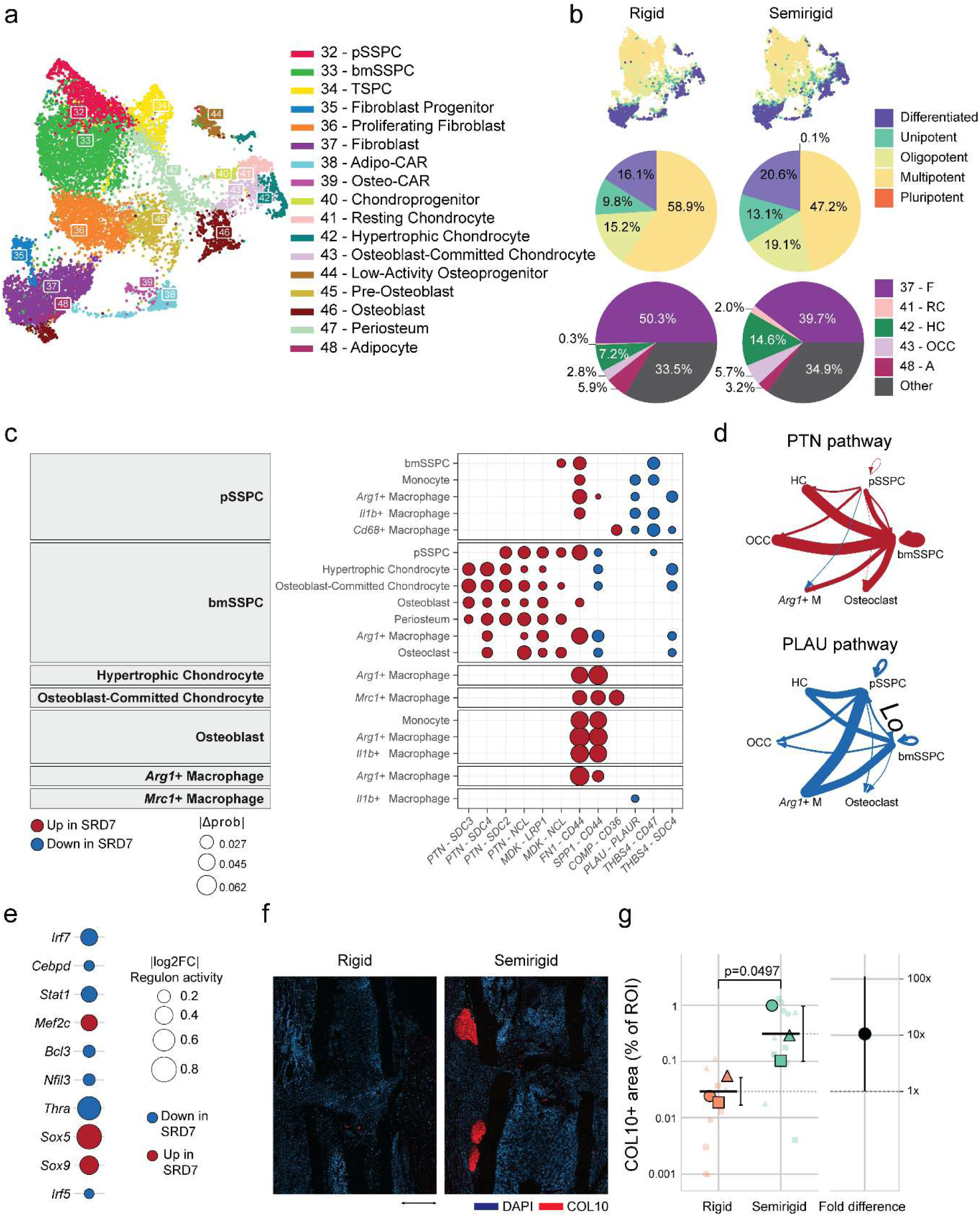
Semirigid fixation is associated with an amplified chondrogenic program by day 7. **a**, UMAP of mesenchymal and skeletal populations identified across the fracture callus, colored by annotated cell type. **b**, Top, UMAPs colored by predicted differentiation potency (CytoTRACE 2) under rigid and semirigid fixation. Middle, proportion of cells in each potency category. Bottom, composition of differentiated populations. **c**, Differential ligand–receptor interactions between skeletal progenitor, chondrocyte, osteogenic and macrophage populations at day 7. Senders are indicated at left, receivers on the vertical axis. Dot color indicates interactions increased (red) or decreased (blue) under semirigid fixation; dot size indicates the absolute difference in communication probability. **d**, Differential communication networks for the PTN (top) and PLAU (bottom) pathways. Arrows run from sender to receiver, width is proportional to the difference in communication probability, and color indicates direction as in **c**. **e**, Differential regulon activity between semirigid and rigid fixation at day 7. **f,** Immunofluorescence of the fracture callus at day 7 under rigid and semirigid fixation, stained for COL10 (red) with DAPI (blue). Scale bar, 500 µm. **g,** COL10+ area as percentage of the region of interest. Small symbols show individual sections (three to four per animal) and large symbols per-animal means, with symbol shape identifying the animal; horizontal lines show geometric means with geometric SD. Right, fold difference (semirigid/rigid) with 95% CI. n = 3 animals per group; two-sided Welch’s t-test on log10-transformed per-animal means. n = 4 to 5 biological replicates per condition and timepoint for all single-cell analyses. SRD7: semirigid day 7, RD7: rigid day 7, pSSPC: Periosteal Skeletal stem and Progenitor Cell, bmSSPC: Bone Marrow Skeletal Stem and Progenitor Cell, TSPC: Tendon Stem and Progenitor Cell, CAR: CXCL12-Abundant Reticular Cell; HC: Hypertrophic Chondrocyte, OCC: Osteoblast-Committed Chondrocyte, RC: Resting Chondrocyte, F: Fibroblast, A: Adipocyte.

Comparison of predicted communication among skeletal progenitor, periosteal and macrophage populations at day 5 showed that matrix-associated signaling toward the macrophage compartment was broadly reduced under semirigid fixation. This includes SPP1–CD44 and SPP1–ITGAV, THBS1–CD36, THBS1–SDC4 and THBS1–CD47, FN1–CD44, and POSTN–ITGAV–ITGB5 (Supplementary Figure 5). Against this reduction, two classes of signal were increased: COMP–SDC4 and COMP–CD47 signaling from periosteum and pSSPCs toward *Mrc1*-expressing, *Il1b*-expressing and *Arg1*-expressing macrophages, and MDK–NCL and MDK–ITGA4–ITGB1 signaling from the same progenitor populations. LGALS9–P4HB signaling from *Mrc1*-expressing macrophages was also increased. The composition of matrix-derived signals directed toward the macrophage compartment therefore differed at day 5, preceding the progenitor growth factor changes observed at day 7.

At day 7, comparison of predicted communication among skeletal progenitor, chondrocyte, osteogenic, and macrophage populations showed a coordinated increase in growth factor signaling from bone marrow-derived skeletal stem and progenitor cells (Figure 3c). PTN signaling through SDC2, SDC3, SDC4 and NCL from bmSSPCs toward hypertrophic and osteoblast-committed chondrocytes, osteoblasts, periosteum, pSSPCs and osteoclasts was increased under semirigid fixation, as was MDK–LRP1 and MDK–NCL signaling from both progenitor populations (Figure 3c–d).

Matrix-associated signaling toward the macrophage compartment was also increased, including FN1–CD44 and SPP1–CD44 from osteoblasts, hypertrophic chondrocytes and osteoblast-committed chondrocytes, and COMP–CD36 from chondrocyte populations toward *Mrc1*-expressing macrophages. In contrast, PLAU–PLAUR signaling from pSSPCs, bmSSPCs and *Mrc1*-expressing macrophages toward monocytes and macrophages was uniformly reduced, as was THBS4–CD47 and THBS4–SDC4 signaling from pSSPCs (Figure 3c–d).

Gene regulatory network analysis on day 7 showed increased *Sox5*, *Sox9* and *Mef2c* regulon activity and reduced *Irf7*, *Stat1* and *Thra* activity under semirigid fixation (Figure 3e). Consistent with a more advanced chondrogenic phase and with the higher proportion of hypertrophic chondrocytes (Figure 3b), COL10+ area within the callus was significantly higher under semirigid fixation, on average 11-fold that of rigid fixation (p = 0.0497; Figure 3f,g).

Taken together, semirigid fixation was associated with a shift toward chondrogenic progenitor states and stage-specific signaling that coincided with increased cartilage formation.

### Mechanosensitive signaling is required for regeneration and pharmacological activation benefits only rigid fixation

To identify a molecular route through which these mechanical differences could act, we examined expression of mechanotransduction-associated genes across the single-cell dataset. *Piezo1* was detected across most populations within the fracture callus, whereas *Trpv4*, *Yap1* and *Ptk2* showed more restricted expression (Figure 4a). *Piezo1* expression shifted between compartments over time, predominating in inflammatory populations including *Arg1*-expressing macrophages and neutrophils at day 1, and in *Mrc1*-expressing macrophages and mesenchymal populations at later timepoints, the same populations in which fixation-dependent signaling differences were observed (Supplementary Figure 6a).

**Figure 4.**
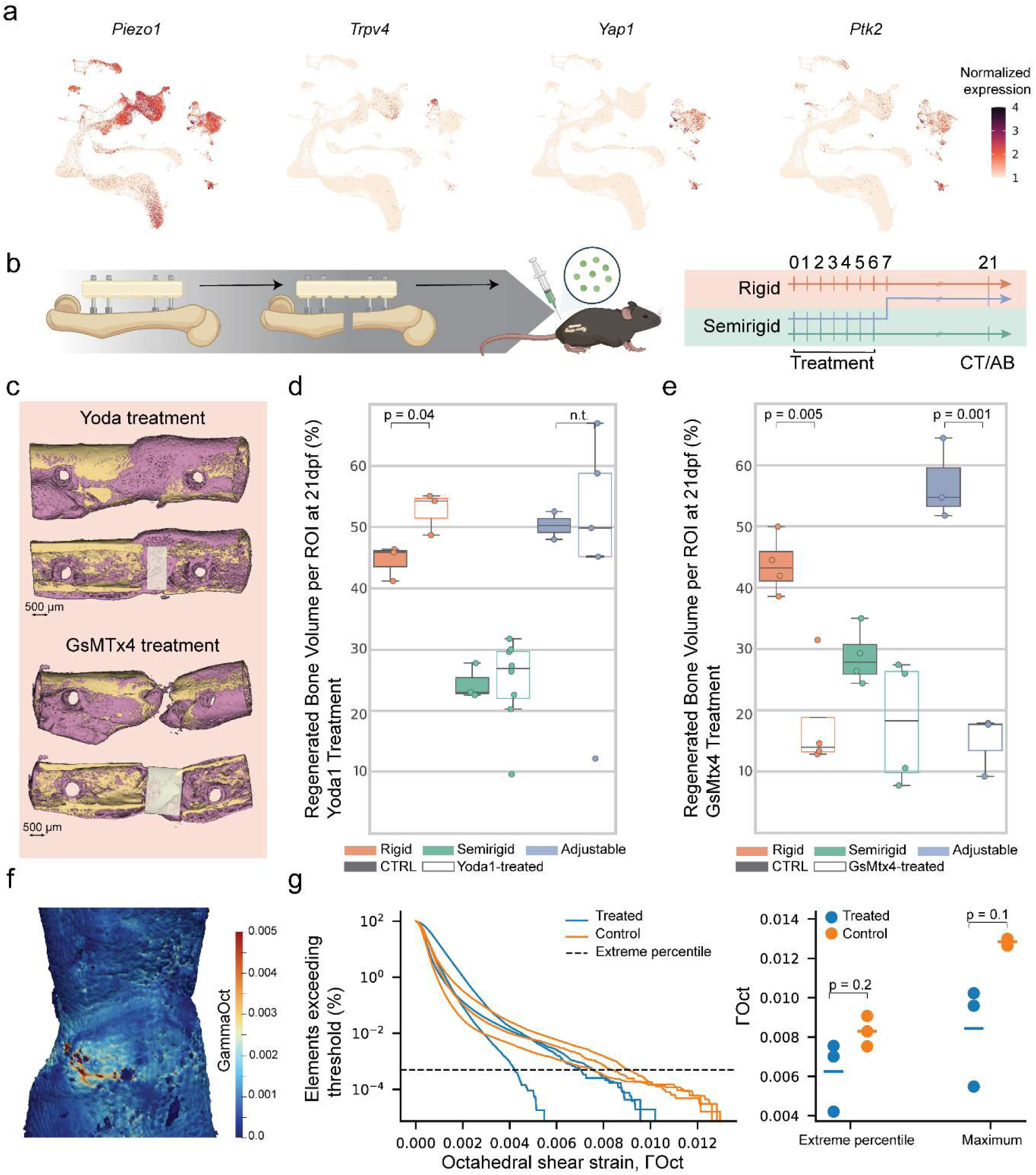
Pharmacological modulation of mechanosensitive channel activity alters fracture repair in a fixation-dependent manner. **a**, Expression of *Piezo1*, *Trpv4*, *Yap1* and *Ptk2* projected onto the integrated UMAP of all cell populations. Color indicates normalized expression. **b**, Schematic of the mouse long bone osteotomy model stabilized by three fixation strategies: rigid fixation, semirigid fixation, and the dynamically adjusted semirigid-to-rigid strategy. Yoda1 or GsMTx4 was administered daily for the first 7 days after osteotomy. Samples were collected on days 1, 3, 5 and 7 for bulk RNA-seq, and on day 21 for CT analysis and Alcian Blue staining. **c**, Representative CT images of calluses at 21 days following Yoda1 (top) and GsMTx4 (bottom) treatment. Cortical bone in yellow, regenerated bone in purple. Scale bar, 500 µm. **d**, Regenerated bone volume per region of interest at 21 days following Yoda1 or vehicle treatment under each fixation condition. Filled boxes, vehicle; open boxes, Yoda1. Boxes show median and interquartile range; points show individual animals. n = 3 and 3 (rigid), 3 and 8 (semirigid), 2 and 5 (dynamically adjusted). Two-sided Welch’s t-test, uncorrected; the dynamically adjusted comparison was not tested owing to the size of the vehicle group (n.t.). **e**, Regenerated bone volume per region of interest at 21 days following GsMTx4 or vehicle treatment under each fixation condition. Filled boxes, vehicle; open boxes, GsMTx4. Boxes show median and interquartile range; points show individual animals. n = 4 and 4 (rigid), 4 and 4 (semirigid), 3 and 3 (dynamically adjusted). Two-sided Welch’s t-test, uncorrected. **f**, Representative distribution of octahedral shear strain (ΓOct) within the region of interest of a healed femur under simulated torsional loading. **g**, Left, exceedance curves showing the proportion of elements within the region of interest experiencing octahedral shear strain above a given threshold, for vehicle-treated and Yoda1-treated specimens under rigid fixation. Dashed line indicates the extreme percentile threshold. Right, extreme percentile and maximum ΓOct values per specimen. n = 3 specimens per group; compared by exact permutation test on group means, for which the smallest attainable two-sided p-value at this sample size is 0.1. Group sizes, means, effect sizes and Holm-adjusted p-values for **d** and **e** are given in Supplementary Tables 3 and 4.

Given the expression of *Piezo1* across immune and stromal populations and its association with cellular changes observed under different mechanical conditions, we hypothesized that modulation of PIEZO1 activity contributes to the effect of the mechanical environment on fracture healing. To test this, mice subjected to rigid, semirigid, or dynamically adjusted semirigid-to-rigid fixation received daily injections during the first 7 days following osteotomy with either Yoda1, a PIEZO1 agonist, or GsMTx4, a non-selective blocker of mechanosensitive cation channels including PIEZO1 (Figure 4b). Among the documented alternative targets of GsMTx4, *Trpc1* and *Trpc6* were detected at low levels within the callus, *Piezo2* expression was largely confined to mesenchymal populations while *Tmem120a* was broadly expressed (Supplementary Figure 6b).

Micro-CT analysis on day 21 demonstrated that pharmacological activation of PIEZO1 with Yoda1 increased regenerated bone volume under rigid fixation, from 44.5 ± 2.9% to 52.7 ± 3.5% (p = 0.036). No difference was observed under semirigid fixation (24.5 ± 2.9% to 24.7 ± 7.2%, p = 0.94). Under dynamically adjusted fixation, values were similar between treated and control animals (46.6 ± 20.9% versus 50.3 ± 3.2%). Yoda1 therefore increased bone formation in the fixation condition providing the least mechanical stimulation, without a corresponding effect under the more compliant conditions (Figure 4c,d, Supplementary Table 3).

Conversely, inhibition of mechanosensitive channel activity with GsMTx4 reduced regenerated bone volume under rigid fixation, from 43.7 ± 4.8% to 18.1 ± 9.0% (p= 0.005), and under dynamically adjusted fixation, from 57.0 ± 6.6% to 14.9 ± 5.0% (p = 0.001). A reduction of the same direction but smaller magnitude was observed under semirigid fixation (28.8 ± 4.6% to 17.9 ± 10.2%, p = 0.12). Inhibition of mechanosensitive channel activity therefore reduced bone formation under all three fixation conditions, with the largest effects in the conditions that showed the greatest bone formation in untreated animals (Figure 4c,e, Supplementary Table 4).

To determine whether the differences in bone bridging affected mechanical competence, torsional loading was simulated using a finite element model of the healed femora from rigidly fixed control and Yoda1-treated animals (Supplementary Figure 7). Octahedral shear strain (Γ_Oct) was used to characterize the local mechanical strain distribution. Γ_Oct was heterogeneous throughout the region of interest, with localized areas of elevated strain (Figure 4f). Exceedance curves showed broadly similar strain distributions, with Yoda1-treated specimens exhibiting a shorter upper tail and lower strains at the highest percentiles (Figure 4g). Maximum Γ_Oct was lower in every treated specimen than in every control specimen (0.0055–0.0102 versus 0.0126–0.0130; Hedges’ g = 1.93). With three specimens per group, an exact permutation test cannot return a p-value below 0.1 (p = 0.1), so this comparison is reported as an effect size rather than as a hypothesis test.

Bulk RNA sequencing (bulk RNA-seq) of fracture calluses confirmed sustained elevation of *Piezo1* transcripts following Yoda1 administration identifying partial overlap between the transcriptional responses to semirigid fixation and PIEZO1 activation. That included increased expression of genes associated with regulatory immune responses and pro-regenerative macrophage states (Supplementary Figure 8).

Together, these findings indicate that PIEZO1 activity contributes to fracture healing and that the effect of PIEZO1 activation depends on the mechanical environment. PIEZO1 activation enhanced bone bridging under rigid fixation, whereas PIEZO1 inhibition impaired bone bridging across all fixation conditions.

### The mechanical environment acts at successive stages of early repair

Taken together, these observations describe a sequence of stage-specific changes rather than a single point of divergence (Figure 5). Within one day of injury, semirigid fixation increased signaling among the resident myeloid populations of the fracture hematoma, including TNF, PPIA and integrin-mediated interactions between mast cells, neutrophils and *Arg1*-expressing macrophages, while reducing chemokine signaling directed toward monocytes and signaling associated with resolution of inflammation. By day 5, the matrix-associated signals reaching the macrophage compartment had changed in composition, with cartilage-associated ligands from periosteal and progenitor populations increasing (Supplementary Figure 5). By day 7, pleiotrophin and midkine signaling from skeletal progenitors toward chondrocyte populations was elevated accompanied by increased *Sox9* regulon activity and more extensive COL10+ cartilage matrix, while plasminogen activator signaling toward myeloid populations was reduced (Figure 3). *Piezo1* was expressed across the populations showing these differences at each stage and shifted between compartments as the responding populations changed.

**Figure 5.**
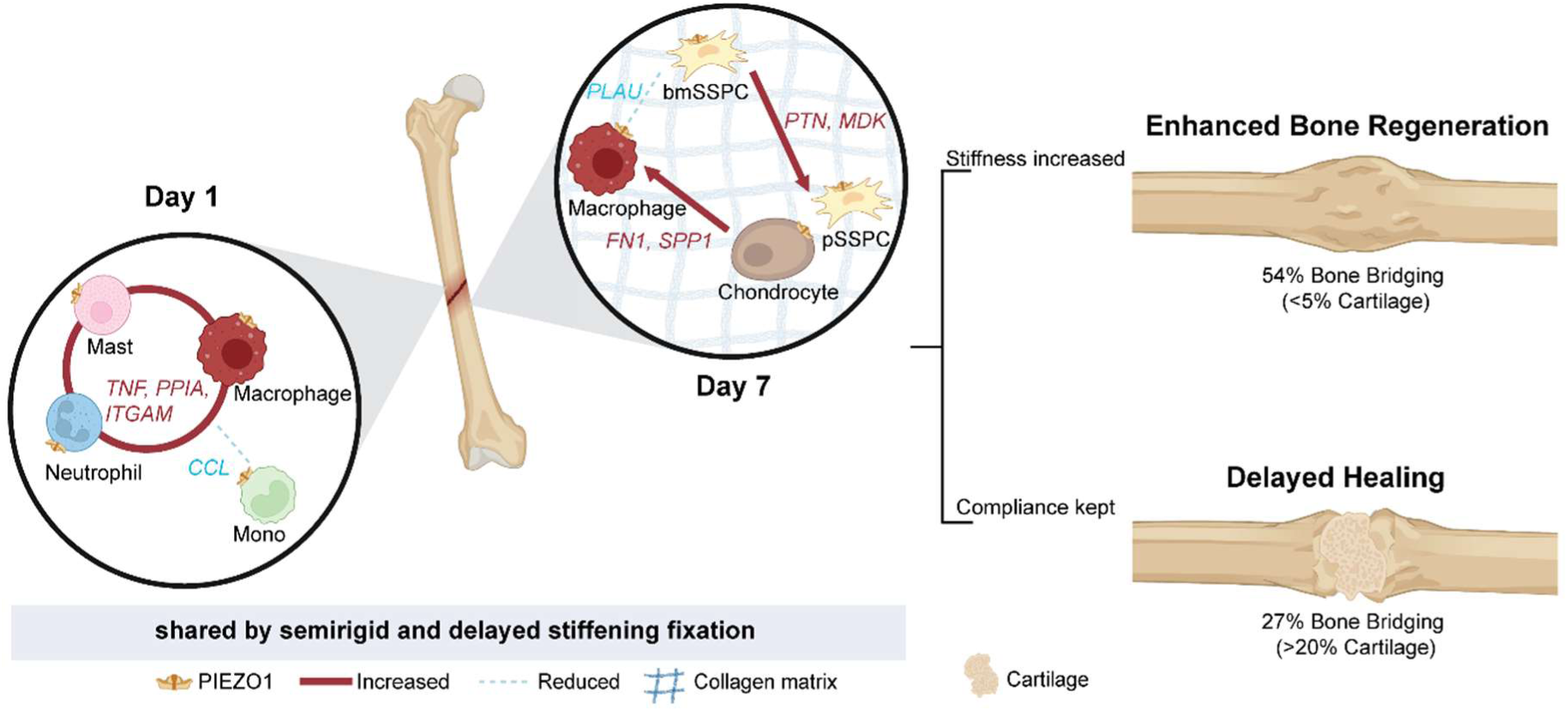
The mechanical environment acts at successive stages of early repair. Schematic summary of the changes in predicted intercellular signaling identified in this study and their relationship to healing outcome. Left and center, magnified views of the fracture callus at 1 and 7 days after osteotomy. At day 1, signaling among mast cells, neutrophils and *Arg1-*expressing macrophages was increased under semirigid fixation, including TNF, PPIA and integrin-mediated interactions, whereas CCL-mediated signaling toward monocytes was reduced. On day 7, pleiotrophin (PTN) and midkine (MDK) signaling from bmSSPCs toward pSSPCs and chondrocytes was increased, as was fibronectin (FN1) and osteopontin (SPP1) signaling from chondrocytes toward macrophages, whereas plasminogen activator (PLAU) signaling from bmSSPCs toward macrophages was reduced. The grid indicates the collagen-rich matrix of the developing callus. *Piezo1* was expressed across the responding populations at both stages and shifted from inflammatory to mesenchymal compartments as healing progressed. Solid red lines indicate increased and dashed blue lines reduced predicted communication under semirigid compared with rigid fixation. Gray bar, the mechanical conditions of the first 7 days are shared by the semirigid and dynamically adjusted groups, which differ only from day 7 onward. Right, healing outcomes on day 21: increasing fixation stiffness at day 7 was followed by 54% bone bridging with less than 5% residual cartilage, whereas maintained compliance was followed by 27% bridging with more than 20% residual cartilage (quantified in Figure 1c).

These changes occurred under mechanical conditions shared by the semirigid and dynamically adjusted groups, which experience the same fixation configuration until day 7 and differ only thereafter. From that point the two strategies diverge: increasing fixation stiffness was followed by the greatest bone bridging and the lowest residual cartilage of the three conditions at day 21, whereas maintaining compliance was followed by persistent cartilage and reduced bridging (Figure 1c, Figure 5).

## Discussion

This study shows that the mechanical environment during the first week after fracture determines how repair proceeds. Fixation stiffness altered intercellular signaling within one day of injury and progressively reshaped the callus toward a chondrogenic program by day 7, and increasing stiffness after this window produced better regeneration than constant rigid fixation. PIEZO1 was expressed across the responding populations and its pharmacological activation improved healing where mechanical stimulation was lowest.

Fixation stiffness during the earliest phase of bone regeneration has long-lasting consequences for repair outcome. Rigid fixation consistently produced successful bone bridging, whereas semirigid fixation delayed healing. Increasing fixation stiffness after the first 7 days produced the best outcome of the three strategies, exceeding constant rigid fixation. The mechanical environment during the initial inflammatory phase therefore does not simply set the pace of repair, it establishes the regenerative trajectory of the callus.

Studies of mechanical stability in fracture healing have historically concentrated on the reparative and remodeling phases^15–17^. Delayed stabilization has been shown to increase cartilage formation, identifying the early fracture environment as a critical window in which mechanical conditions steer tissue fate^18^. Unlike previous models that delayed stabilization entirely, our approach maintained continuous fixation while selectively altering fixation stiffness, demonstrating that repair is regulated not only by the presence or absence of mechanical stability, but by the magnitude and temporal evolution of mechanical signals. Constant rigid fixation supported bridging but was outperformed by a strategy in which stabilization was increased only after the first week, whereas maintaining reduced stiffness throughout delayed repair. The three constructs differ in absolute stiffness, so these results establish that a strategy of early compliance followed by increased stiffness outperforms constant fixation at either stiffness, rather than isolating timing as an independent variable.

The earliest detectable differences among fixation conditions in the single-cell data were directional rather than quantitative. Under semirigid fixation, signaling among the resident myeloid populations of the hematoma was maintained or increased while signaling toward monocytes was reduced, despite monocytes being more abundant, a dissociation indicating that the difference lies in how myeloid cells are engaged rather than how many arrive. Two observations point in the same direction. TNF was redirected toward TNFRSF1B, which lacks a death domain and preferentially activates survival-associated signaling^19^, consistent with the increased NF-κB regulon activity observed in the same samples by an independent analysis. Engagement of the formyl peptide receptors shifted in ligand rather than in magnitude: annexin A1, which promotes neutrophil apoptosis and macrophage efferocytosis through FPR2^20^, was reduced, while cathepsin G, a chemotactic FPR1 agonist released from granules^21^, was increased. Semirigid fixation therefore appears to sustain and retain the initial myeloid response rather than amplify it, while reducing the signals that would normally initiate its resolution.

The shift from the inflammatory to the chondrogenic phase was already apparent on day 5, when signaling from periosteal and progenitor populations toward macrophages changed in composition rather than in magnitude. Osteopontin, thrombospondin-1, fibronectin and periostin, all associated with bone matrix, were reduced in the semirigid condition, while cartilage oligomeric matrix protein and midkine from the same populations were increased. Because macrophages read the surrounding matrix through CD44, CD47 and integrin receptors^22,23^, this represents a change in the matrix context to which the immune compartment is exposed and provides a plausible link between the altered inflammatory signaling at day 1 and the amplified chondrogenic program at day 7. Progenitor-derived midkine signaling toward macrophages has also been reported in the healing periosteum by spatial transcriptomics^24^, supporting this axis as a feature of normal repair that is modulated by the mechanical environment.

By day 7 the effects of fixation stiffness had shifted to the skeletal compartment. Pleiotrophin and midkine signaling from bone marrow-derived progenitors toward chondrocyte and osteogenic populations was increased, alongside elevated *Sox5* and *Sox9* regulon activity and a larger COL10+ hypertrophic cartilage area. Pleiotrophin and midkine are both heparin-binding growth factors implicated in chondrogenic differentiation, and midkine-deficient mice show delayed chondrogenesis during early fracture healing^25^. Because day 7 corresponds to the soft callus stage, these observations indicate that the chondrogenic phase proceeds more strongly under semirigid fixation rather than being diverted.

Signaling toward the macrophage compartment showed a coordinated shift under semirigid fixation. Fibronectin and osteopontin signaling through CD44 from osteoblasts and chondrocytes were increased, while plasminogen activator signaling toward the same populations was uniformly reduced. CD44 ligation on macrophages promotes retention within the matrix, phagocytic clearance of apoptotic cells, activation of pro-MMP9 and macrophage fusion^22^. These are the functions required for cartilage resorption at this stage. Plasmin generation, in contrast, is required for matrix degradation and osteoclast-mediated remodeling of fracture cartilage^26^. Under semirigid fixation, the callus therefore enters the second week with matrix signals directing resorption but reduced proteolytic capacity to carry it out, consistent with the cartilage that persists to day 21.

Where this trajectory leads to, if flexibility is maintained throughout healing, is illustrated by a recent study in which high interfragmentary strain was sustained throughout healing^27^. Fractures stabilized under high strain developed larger but biomechanically inferior calluses than low-strain controls, resembling hypertrophic nonunion, and this deficit was associated with altered tissue organization and a fibrotic, immune-associated transcriptional state rather than an impaired capacity for mineralization. Early flexibility is therefore not harmful in itself but becomes so if it persists. It appears to support the early inflammatory and chondrogenic phases, while subsequent stiffening is needed for progression to organized bone formation.

PIEZO1 may contribute to these responses in two ways as suggested by our data. It was expressed across the populations showing fixation-dependent differences and shifted between compartments as those populations changed, being detected predominantly in inflammatory populations at day 1 and in pro-regenerative macrophage and mesenchymal populations later in healing. This distribution is consistent with a role as a mediator translating mechanical conditions into cellular responses at successive stages of repair, and semirigid fixation increased NF-κB regulon activity, a pathway previously linked to PIEZO1-mediated calcium signaling^28,29^. That pathway can also be activated independently of PIEZO1, so the expression and regulon data are compatible with such a role without establishing it.

The pharmacological experiments further support the notion that mechanical stimulation during the early healing window promotes bone regeneration, and that cellular sensitivity to mechanical cues can modulate this response. Yoda1 treatment significantly improved healing under rigid fixation but produced no additional benefit under semirigid or dynamically adjusted fixation, whereas GsMTx4 treatment impaired repair across both the rigid and adjustable condition, indicating that mechanosensitive signaling contributes to successful healing. Finite element analysis of the healed calluses further showed lower extreme octahedral shear strains following Yoda1 treatment, consistent with the increased bone bridging, although the small number of specimens limits this to supporting evidence. Whether PIEZO1 also mediates the endogenous response to fixation stiffness is a separate question these experiments do not resolve, and one that conditional deletion in defined cell populations would be required to address. Pharmacological modulation cannot fully exclude effects on additional mechanosensitive channels, particularly for GsMTx4, which also inhibits TRPC channels, PIEZO2 and TACAN (TMEM120A)^30,31^. Within the fracture callus, *Trpc1* and *Trpc6* were detected at low levels, and *Piezo2* expression was confined to mesenchymal populations, making a substantial contribution from these channels unlikely, although *Tmem120a* was broadly expressed and a contribution cannot be excluded. Nevertheless, irrespective of whether mechanical stimulation was increased through more compliant fixation or cellular mechanosensitivity was pharmacologically enhanced, interventions that increased mechanical input or mechanosensitive responsiveness during the early healing window were associated with improved regeneration.

At the transcriptional level, systemic Yoda1 administration produced sustained elevation of *Piezo1* transcripts within the cells of the defects throughout the first week of healing, demonstrating a transcriptional response to treatment within the healing tissue, although indirect effects of systemic PIEZO1 activation in other tissues cannot be excluded. Gene set enrichment analysis identified calcium-associated pathways early after treatment, in keeping with PIEZO1-mediated calcium signaling, alongside immune regulatory and cellular activation pathways, implicating PIEZO1 activation in both mechanosensitive signaling and the inflammatory state of the healing callus.

Genetic studies place PIEZO1 in the skeletal compartment at later stages of repair. PIEZO1 in chondrocytes is required for normal healing, its loss driving inflammatory and oxidative stress responses within chondrocytes^32^, while deletion under the collagen 10 promoter leaves hypertrophic chondrocyte apoptosis and transdifferentiation intact but derepresses Receptor Activator of Nuclear Factor-κB Ligand (RANKL) and reduces osteoprotegerin expression, increasing osteoclast numbers and impairing callus maturation^33^. Together with the shift in *Piezo1* expression from inflammatory to mesenchymal populations in our dataset, this suggests that PIEZO1 acts at successive stages through different compartments rather than exerting a single effect on inflammation. Our day 7 data point to the interface between hypertrophic cartilage and the resorptive compartment through a different effector, with plasminogen activator signaling toward macrophages reduced under semirigid fixation. Our pharmacological data are consistent with, but do not by themselves establish, a comparable endogenous role during fixation-dependent repair.

Our study does have several limitations. Construct stiffness differed slightly between the adjustable and non-adjustable fixator systems supplied by the manufacturer (Supplementary Table 1), such that the dynamically adjusted group was not exactly equivalent to the semirigid and rigid groups in the respective phases of regeneration. Nevertheless, the adjustable fixator started at a slightly lower stiffness than the non-adjustable semirigid system and did not reach the stiffness of the rigid system after adjustment. Despite this more limited increase in stiffness, delayed stiffening produced greater bone regeneration than constant rigid fixation, supporting the importance of the timing of stiffness rather than absolute stiffness alone. Pharmacological modulation cannot fully exclude effects on additional mechanosensitive channels, as discussed above. The Yoda1-treated samples were sequenced on a different platform from the untreated rigid and semirigid samples, without contemporaneous controls, and the two cohorts also differed in osteotomy technique, housing and sample pooling, so treatment effects cannot be fully separated from cohort and platform effects in that comparison. The semirigid versus rigid contrast was performed entirely within a single sequencing batch. Finally, all experiments used male mice; sex-dependent differences have been reported in both the inflammatory response to fracture and in PIEZO1-mediated mechanotransduction and should be examined in future work. Together, these considerations define important boundaries for interpretation while leaving the central finding that the timing of mechanical stiffening influences the regenerative outcome supported by the independent fixation and pharmacological experiments.

A complete account of this study will require analysis of the second week of healing. Our profiling was designed to capture the early response up to day 7, when the adjustable semirigid-to-rigid fixation was switched to rigid fixation. The events that follow stiffening, including whether the amplified chondrogenic program subsequently converts toward bone formation or persists as cartilage, therefore remain uncharacterized. Profiling the adjustable group after stiffening could determine whether increased stiffness restores the proteolytic signaling reduced under semirigid fixation at day 7 or instead acts more directly on chondrocytes. Extending outcome measures beyond day 21 would further establish whether the advantage of delayed stiffening persists through remodeling. More broadly, varying the timing of the stiffness transition will be important to determine whether the mechanosensitive window identified here can be further optimized.

Collectively, fixation stiffness regulates fracture repair by coordinating the mechanical environment with immune responses and tissue regeneration. The early mechanical environment acts not only as a structural constraint but as a biological signal that shapes the inflammatory microenvironment and directs the subsequent regenerative trajectory of the callus, with PIEZO1 emerging as a candidate mediator. Successful regeneration therefore requires not simply maximal stability or maximal mechanosensory activation, but temporal coordination between mechanical input and cellular state. This principle has implications for the design of fixation strategies and regenerative biomaterials, where mechanical cues may need to evolve with the changing requirements of the healing tissue.

## Materials and methods

### Mice

Male C57BL/6 mice were obtained from Janvier Labs. Mice used for the single-cell and untreated bulk RNA sequencing experiments were housed in a conventional facility; mice used for all experiments analyzed on day 21, including the pharmacological experiments, were housed under specific pathogen-free conditions. All mice were kept on a 12 h light/dark cycle with ad libitum access to food and water. Ten- to twelve-week-old mice were used for all experiments. All animal procedures were approved by the KU Leuven Ethical Committee (p081/2023, p051/2025) and conducted in accordance with institutional and national guidelines. Animals were randomly assigned to experimental groups. Investigators were not blinded during surgery, treatment and sample collection, as the fixator type was visible. Micro-CT scans were acquired by a technician unaware of group allocation. All subsequent image analyses, including micro-CT, histology and immunofluorescence quantification, were performed on samples identified only by mouse ID, and group allocation was assigned after analysis was completed.

### Femoral Defect Surgery

Mice were anesthetized via intraperitoneal injection of ketamine (80 mg/kg) and xylazine (5 mg/kg) and received buprenorphine (0.1 mg/kg) for perioperative analgesia. Animals were maintained on a 37 °C heating pad throughout the procedure. The right hindlimb was shaved and disinfected, followed by a skin and fascia incision and blunt muscle splitting to expose the femur. A unilateral external fixator (RISystem) was applied using four screws. A standardized mid-diaphyseal osteotomy was performed using a 0.66 mm wire saw. For the single-cell and untreated bulk RNA sequencing experiments, the osteotomy was instead made with two cuts using a 0.22 mm wire saw, producing a gap of approximately 0.5 to 1 mm. All animals analyzed on day 21, including the pharmacological experiments, received 0.66 mm osteotomy. The wound was closed with sutures, and mice received two additional buprenorphine injections during the first 24 h post-surgery. Mice were euthanized at 1, 3, 5, 7, or 21 days post-fracture.

### Drug administration

Yoda1 stock solutions (50 mM, SML1558, Sigma-Aldrich) were prepared in DMSO and stored at −80 °C. Immediately before use, aliquots were diluted in 5% ethanol in 0.9% NaCl. Yoda1 was administered intraperitoneally at 5 µmol/kg/day, with the injection volume adjusted to body weight (approximately 100 µl), for 7 consecutive days starting on the day of osteotomy. GsMTx4 stock solutions (5 mM, HY-P1410, MedChemExpress) were prepared in DMSO and stored at −80 °C. Immediately before use, aliquots were diluted in 0.9% NaCl. GsMTx4 was administered intraperitoneally at 1 mg/kg/day, with the injection volume adjusted to body weight (approximately 100 µl), following the same schedule as Yoda1.

### Instron testing for fixator stiffness

The RISystem external fixator was assembled on a PEEK rod (Ø 2 mm) with predrilled holes and mounted in a collet in an Instron 5886 universal testing machine (Instron, USA) fitted with a ±10 N load cell. The collet was aligned with a flat indenter in the loading head, and the crosshead lowered until a compressive contact force of 0.1–0.15 N was reached against the proximal rod of the construct. Three loading cycles from 0 to 60 µm displacement were applied, with force and displacement recorded continuously. Data were processed in MATLAB (v26.1), and construct stiffness was taken as the slope of a linear regression fitted to the third loading cycle over 0–0.06 mm of displacement. One construct was tested per fixator configuration, so the reported stiffness values characterize the nominal mechanical behavior of each system rather than variability between individual constructs. Construct stiffness differed between the adjustable and non-adjustable fixator systems supplied by the manufacturer, such that the adjustable construct was less stiff than the non-adjustable construct at both the 50% and 100% configurations (4.33 versus 6.51 N/mm and 11.41 versus 20.38 N/mm, respectively; Supplementary Table 1). The dynamically adjusted group therefore experienced a more compliant environment than the semirigid group during the first 7 days, and a less rigid environment than the rigid group thereafter.

### Sampling and preparation

For micro-CT, histology and immunofluorescence, femurs were harvested following euthanasia at 1, 3, 5, 7 and 21 days post-fracture, fixed overnight in 4% paraformaldehyde (PFA) in phosphate-buffered saline (PBS) at 4 °C, and processed for micro-CT acquisition, histology or immunofluorescence as described below.

### Micro-CT acquisition

Samples were scanned for mineralized tissue analysis using a Phoenix Nanotom M micro-CT system (Baker Hughes) in FastCT mode, in which projections are acquired during continuous rotation of the sample. Samples were aligned along the longitudinal axis of the femur. Scanning parameters included a diamond – tungsten target, 60 kV voltage, 170 µA current, 500 ms exposure time, 1800 projections over 360° without frame averaging, and a 0.2 mm aluminum filter, resulting in a voxel size of 5 µm.

### Micro-CT analysis

Raw projection data were reconstructed using CT Pro 3D software (Nikon Metrology) and imported into 3D Slicer (version 5.6) for analysis. A cylindrical region of interest (ROI) was defined in line with the cortical bone, spanning the osteotomy gap between the cut ends, and debris was removed using segmentation masks to isolate the femur and callus. Mineralized tissue was identified using the moments thresholding algorithm. Threshold values were determined based on histogram inspection. Newly formed and cortical bone were differentiated by adjusting threshold values. Bone volumes within the ROI were calculated in mm³, and the volume of newly formed bone was expressed as a percentage of the ROI volume (bridged bone percentage).

### Micro-CT-based in silico torsion testing

For each sample, the micro-CT volume was cropped to the bone segment between the two outer fixation pins and segmented using the intensity threshold described above. Small, isolated components were removed, and the resulting volumes were aligned to the longitudinal axis of the bone and downsampled fourfold while preserving the physical dimensions. A four-node tetrahedral finite element mesh (C3D4) was generated from the segmented volume. Voxel intensity was converted to apparent mineral density using a quadratic calibration function derived from borosilicate (2.23 g/cm³) and ceramic (3.85 g/cm³) reference spheres included in the micro-CT scans. Young’s modulus was calculated from apparent mineral density using the power-law relationship E = a · ρ^b^ where ρ is in g/cm³ and E in MPa. The exponent (b = 2) was taken from a previously described finite element model of the mouse femur^34^, and the coefficient was set to a = 1.4 × 10³ so that cortical bone elements reached Young’s modulus values consistent with those measured by indentation in femoral cortical bone of C57BL/6 mice^35,36^. A Poisson’s ratio of 0.4 was assigned to all elements. Elements with Young’s modulus values within 1% were grouped into common material sets, and mesh quality and the spatial distributions of density and Young’s modulus were visually verified before finite element analysis.

The generated models were imported into Abaqus 2020 and subjected to a static torsional loading protocol. The proximal fixation plate was coupled to a reference point, and a 15 Nmm torsional moment was applied, while the distal fixation plate was constrained with encastre. The applied torsional moment was based on ultimate torque values reported for mouse femora in experimental torsion studies.^37^ Following simulation, element-level strains were extracted from the final analysis frame. Octahedral shear strain was calculated from elements within a predefined longitudinal ROI centered on the fracture site, spanning 1.3 mm along the x-axis. The resulting strain distributions were summarized by their mean, standard deviation, maximum, and the 99.9995th percentile. Because most elements within the region of interest experience little strain under the applied loading, mean values are insensitive to differences between specimens, and the upper tail of the distribution was used instead. The maximum reflects a single element within meshes of several million; the 99.9995th percentile was therefore included as a more stable measure of the upper tail.

### Histology

After micro-CT acquisition, samples were washed in PBS, decalcified in 0.5 M EDTA in PBS for 10 days at room temperature with gentle agitation, dehydrated, embedded in paraffin and sectioned at 5 µm. Paraffin sections were deparaffinized and rehydrated prior to staining. Hematoxylin and eosin (HCE) staining was performed following standard protocols. Alcian Blue staining was carried out using a 1% Alcian Blue solution, followed by counterstaining with nuclear fast red. Slides were scanned using the Zeiss Axio Scan.Z1 slide scanner. For Alcian Blue quantification, regions of interest were manually defined between the fracture ends using Fiji. Default thresholding was applied uniformly across samples. Cartilage area was quantified in three to five sections per animal and averaged to give one value per animal, with the individual animal as the experimental unit.

### Immunofluorescence

Paraffin sections were deparaffinized, rehydrated and subjected to antigen retrieval in P6 buffer (S1699, Agilent) at 98 °C for 20 min. For COL10 staining, this was followed by enzymatic retrieval with pepsin (0.5 mg/ml in 0.01 M HCl) for 15 min at room temperature. Sections were blocked for 30 min in 15% donkey serum and 15% goat serum in PBS containing 0.1% Tween-20 and 0.3% Triton X-100 and incubated overnight at 4 °C with primary antibodies against LY6G/C (1:50, 14-5931-82, Thermo Fisher Scientific), MMP9 (1:50, 10375-2-AP, Proteintech), CCR2 (1:50, ab273050, Abcam) and COL10 (1:50, ab260040, Abcam). Sections were then incubated for 2 hours at room temperature with donkey anti-rabbit IgG Alexa Fluor 647 (1:500, A-31573, Thermo Fisher Scientific) and goat anti-rat IgG Alexa Fluor 488 (1:500, A-11006, Thermo Fisher Scientific). For double stainings including the Alexa Fluor 488 channel, autofluorescence was quenched using the Vector TrueVIEW Autofluorescence Quenching Kit (SP-8400, Vector Laboratories). Nuclei were counterstained with DAPI. Staining specificity was confirmed during protocol optimization using sections incubated without primary antibody. For image acquisition, a Zeiss Axioscan Z.1 slidescanner was used equipped with a Hamamatsu Orca Flash 4.0 V3 camera in combination with a Plan-Apochromat 20×/0.8 M27 objective (NA 0.8). The setup was controlled by ZEN blue (v3.13, Carl Zeiss Microscopy GmbH). The sampling rate was 325 nm/pixel. DAPI was excited with a 385 nm LED module and collected with a 412–438 nm emission filter, Alexa Fluor 488 was excited with a 475 nm LED module and collected with a 500–550 nm emission filter, and Alexa Fluor 647 was excited with a 630 nm LED module and collected with a 662–756 nm emission filter. For post-processing image analysis, ZEN blue (v3.13, Carl Zeiss Microscopy GmbH) was used.

### Immunofluorescence cell quantification

LY6G/C/MMP9 and LY6G/C/CCR2 double stainings were quantified at day 1 using a custom Python pipeline based on napari (v0.4.19), scikit-image (v0.23.2), SimpleITK (v2.3.1) and Cellpose (v3.1.1.3), with NumPy v1.26.4. Sections were exported from the slide scanner at full resolution as RGB TIFF files (pixel size 0.325 µm). Rectangular ROIs were placed in the proximal and distal regions of two sections per animal, giving four ROIs per animal (n = 4 rigid and 3 semirigid animals). Each channel was normalized, denoised with a median filter, adjusted for contrast and gamma, smoothed with a Gaussian filter (σ = 0.5 pixels) and enhanced by histogram equalization with clipping. Nuclei were segmented from the DAPI channel with Cellpose (expected nuclear diameter 5 µm) and filtered by size and roundness. Marker channels were binarized at a multiple of the image-specific Otsu threshold. The multiplier (1-2) was adjusted per image by visual inspection of the thresholded output to match the visible staining pattern, without knowledge of group allocation. Marker-positive pixels were assigned to the nearest nucleus using a watershed algorithm, with segmented nuclei as seeds, to define cell regions (expected cell diameter 7 µm). LY6G/C+, MMP9+ or CCR2+, and double-positive cells were expressed as a percentage of all segmented nuclei within each ROI. Segmentation and classification were verified visually using overlay images for each ROI. Percentages were averaged across the two sections within each animal and region, giving one value per animal for the proximal and one for the distal region.

### COL10 immunofluorescence quantification

COL10 immunofluorescence was quantified in Fiji (ImageJ 1.54p) on three to four sections per animal from the defect region between the inner pins (n = 3 animals per group, day 7). Images were exported from the slide scanner at 50% resolution, giving a final pixel size of 0.65 µm. After splitting the color channels, the 8-bit COL10 channel was thresholded at a single fixed intensity value (100 of 255) for all images. This value was set to exclude signal in adjacent skeletal muscle on the same sections, where COL10 is not expected, and approximated the automated (Default) threshold on strongly stained sections. COL10+ area was expressed as a percentage of a region of interest covering the defect between the fracture ends (10.0–11.0 mm²). The same ROI was repositioned in each section and adjusted where needed to match the defect. Values were averaged per animal, with the individual animal as the experimental unit.

### Statistical analysis

The experimental unit throughout was the individual animal. Two sets of *in vivo* experiments were analyzed separately. In the fixation experiment, regenerated bone volume per region of interest at 21 days was compared between rigid, semirigid and dynamically adjusted fixation by one-way ANOVA followed by Tukey’s honestly significant difference test (n = 4, 4 and 3 animals); reported p-values are Tukey-adjusted. Cartilage area was quantified on three to five sections per animal and averaged to give one value per animal, and fixation conditions were compared by Welch ANOVA, which does not assume equal variance between groups, followed by pairwise two-sided Welch’s t-tests with Holm correction for three comparisons (n = 3 animals per group).

The pharmacological experiments comprised three independent cohorts, one per fixation condition. Within each cohort, treated animals were compared with vehicle-treated controls of the same fixation condition, giving one pre-planned comparison per cohort. Because these comparisons address independent questions in independent cohorts, they were analyzed separately by two-sided Welch’s t-test and p-values are reported uncorrected; the number of comparisons performed is stated and Holm-adjusted values are given in Supplementary Tables 3 and 4. Effect sizes are reported as Hedges’ g. Groups comprising fewer than three animals were reported descriptively and not tested; this applied to the dynamically adjusted vehicle group of the Yoda1 experiment (n = 2). Exact group sizes are listed in Supplementary Tables 3 and 4.

Immunofluorescence cell percentages at day 1 were compared between rigid and semirigid fixation separately for the proximal and distal regions by Welch’s t-test, with Holm correction for the two regions within each staining and marker (n = 4 rigid and 3 semirigid animals). The double-positive populations were considered the primary measures, as they identify inflammatory monocytes (CCR2+LY6G/C+) and neutrophils (MMP9+LY6G/C+); single-marker percentages are reported as supporting data.

COL10+ area at day 7 was compared between rigid and semirigid fixation by two-sided Welch’s t-test on log10-transformed per-animal means, because values were right-skewed and variance differed between groups (n = 3 animals per group); the effect is reported as the ratio of geometric means with 95% confidence interval.

Finite element outcomes were compared between Yoda1-treated and control specimens using an exact permutation test on group means (n = 3 per group). With three specimens per group the smallest attainable two-sided p-value is 0.1, so effect sizes and group separation are reported alongside p-values.

### Single-cell isolation

Hematomas, calluses, and uninjured control bone tissue were microdissected and digested in high glucose Dulbecco’s modified Eagle’s medium (DMEM) with GlutaMAX (31966-021, Thermo Fisher Scientific) containing 3 mg/mL collagenase II (17101-015, Thermo Fisher Scientific), 4 mg/mL dispase (17105-041, Thermo Fisher Scientific), and 1% antibiotic-antimycotics (15240062, Thermo Fisher Scientific) for 30 min at 37 °C with rotation. Digestion was repeated twice. Cells were resuspended in high-glucose DMEM with GlutaMAX, filtered through a 70 µm mesh, and subjected to red blood cell lysis using ACK lysis buffer (A1049201, Thermo Fisher Scientific) for 1 min at room temperature. Cells were washed and resuspended in PBS containing 0.04% BSA. Cell viability exceeded 90% for all samples. Group sizes were based on prior work and pilot data.

### RNA extraction and bulk RNA-seq

Three biological replicates were sequenced by condition and timepoint (42 libraries). For untreated samples, each replicate consisted of callus tissue pooled from two mice; Yoda1-treated replicates were from individual mice. To keep sample preparation identical to the single-cell experiments, calluses were first dissociated into single-cell suspensions as described above, and total RNA was isolated from the cell suspension using the RNeasy Mini Kit (74104, Qiagen) according to the manufacturer’s instructions. RNA concentration and purity were assessed from A260/280 and A260/230 ratios. Libraries were prepared with ribosomal RNA depletion at the KU Leuven Genomics Core and sequenced as 2 × 151 bp paired-end reads with dual 8-bp indices on an Illumina HiSeq 4000 (untreated rigid and semirigid samples) or an Illumina NovaSeq X Plus with a 10B flow cell (Yoda1-treated samples). Reads from both sequencing runs were processed together with an identical pipeline: adapter and low-quality sequences were trimmed and read quality was assessed with fastp, with quality additionally inspected with FastQC; trimmed reads were aligned to the GRCm39 (mm39) mouse reference genome with STAR, which also generated gene-level read counts (--quantMode GeneCounts); and alignments were sorted and indexed with SAMtools (version 1.18). One untreated rigid day 5 sample was a clear outlier on principal component analysis, separating from all other samples, and was excluded, leaving two replicates for that condition and three for all others. Differential gene expression analysis was performed using DESeq2, and gene set enrichment analysis using the GSEA preranked approach with genes ranked by log2 fold change and curated mouse gene sets from the Molecular Signatures Database (MSigDB). Because contemporaneous untreated controls were not sequenced alongside the Yoda1-treated samples, untreated replicates were pooled from two mice whereas treated replicates were not, and the two cohorts differed in osteotomy technique and housing, the comparison between Yoda1-treated and untreated rigid fractures is confounded with sequencing platform, sample pooling and cohort.

### scRNA-seq sequencing and data preprocessing

Five biological replicates were sequenced by condition and timepoint. Single-cell RNA-seq libraries were generated using the Chromium Single Cell Next GEM 5′ V2 Library & TCR/BCR Immune Profiling Kit (10x Genomics) and sequenced on an Illumina NovaSeq 6000 using an S4 flow cell (v1 chemistry). Data was processed using Cell Ranger multi (version 8.0.1) with alignment to the mm10 (GRCm38) reference genome. Quality control was performed using the Scater package^38^. Cells were filtered based on total UMI counts, number of detected genes, and mitochondrial read fraction, excluding cells deviating more than three median absolute deviations from the median. Low-abundance genes and duplicate entries were removed. Libraries with insufficient sequencing depth, with cell composition dominated by red blood cells, or that were clear outliers on principal component analysis were excluded, leaving four to five replicates per condition for analysis (37 of 45 libraries).

### Filtering and clustering using Seurat

Filtered datasets were imported into Seurat v4^39^ and processed per timepoint. Normalization was performed using LogNormalize with a scale factor of 1E4. Data were scaled using all genes, and principal component analysis was conducted using the top 2000 highly variable genes selected by the vst method. The number of PCs used for clustering was determined based on elbow plots. Clustering was performed using the Louvain algorithm with dataset-specific resolution parameters. Cell populations were annotated based on the expression of canonical marker genes identified with FindMarkers, cross-referenced with published bone and immune cell atlases. Selected marker gene expression across all annotated populations is shown in Supplementary Figure 1. Contralateral uninjured controls and days 1, 3, 5, and 7 datasets were merged for integrated clustering and annotation.

### Compositional Analysis using scCODA

Cell-type compositional differences for each timepoint between rigid and semirigid fixation were analyzed using scCODA (v0.1.9)^40^, employing a hierarchical Dirichlet– Multinomial model. Rigid fixation was selected as the reference condition based on its role as the standard stabilization strategy. Automatic cell-type reference selection was used following established protocols. Effects were considered credible at a false discovery rate of 0.2.

### Differentiation state analysis using CytoTRACE 2

Cellular differentiation states were assessed using CytoTRACE 2 (v1.1.0)^41^ with default parameters. CytoTRACE scores were visualized on UMAP embeddings and as pie charts summarizing cluster-level differentiation states.

### Single-cell regulatory network inference using SCENIC

Transcription factor regulatory networks were inferred using SCENIC (v1.3.1)^42^ with AUCell (v1.16.0), RcisTarget (v1.14), and GRNboost, alongside the appropriate motif databases. Regulon activity scores were integrated into Seurat objects and used for feature visualization and lineage-level analyses.

### Differential Cell-Cell Communication analysis with CellChat

Cell-cell communication analysis was performed using the CellChat (v2)^43^ package following the published single-dataset and comparative multi-dataset analysis workflows (https://github.com/jinworks/CellChat). Communication probabilities were inferred independently for each experimental condition. Differential cell-cell communication between conditions was assessed by calculating the change in communication probability (Δ probability; Condition 2 − Condition 1) for each ligand-receptor interaction. Interactions were retained if they were significant in at least one condition (p < 0.05 in either condition) and if either the ligand or receptor was identified as differentially expressed between conditions (ligand or receptor p < 0.05).

## Data Availability

Single Cell and Bulk RNA-seq datasets will be made publicly available on Gene Expression Omnibus upon acceptance of the manuscript.

## Acknowledgements

We would like to thank Karen Moermans for technical support with immunofluorescence optimization and scanning, Marina Maréchal for expertise and support with ethical applications, and Janne Vleminckx for technical assistance throughout the project. The authors gratefully acknowledge the VIB BioImaging Core for their support & assistance in image acquisition. Parts of the figures were created using BioRender. Ivanets, M. (2026) https://BioRender.com/k0y962v. Ivanets, M. (2026), https://BioRender.com/215jpho. Ivanets, M. (2026) https://BioRender.com/zpj2tva. Research was funded by the Research Foundation Flanders (FWO Vlaanderen) G.N.: 12C5923N, TV: 1170726N, mERCATOR: 3M220726,; by the European Union’s Horizon Europe Framework Program (HEU/2022-2027) ERC INSTant CARMA (101088919), as well as by the special research fund of the KU Leuven (C24/17/077). The funders had no role in study design, data collection and analysis, the decision to publish, or the preparation of the manuscript. Part of computational resources and services used in this work were provided by the VSC (Flemish Supercomputer Center), funded by the Research Foundation Flanders (FWO) and the Flemish Government – department EWI. This work is part of Prometheus, the KU Leuven RCD division for skeletal tissue engineering (http://www.kuleuven.be/prometheus).

