## Supplementary Information for "An early mechanosensitive window in bone fracture healing shapes long-term repair"

**Supplementary Table 1. Mechanical characterization of the external fixator systems.** Force–displacement curves and construct stiffness for the adjustable and non-adjustable fixator systems in the 100% and 50% side-part configurations, measured under axial compression. The non-adjustable system was used for the rigid (100%, 20.38 N/mm) and semirigid (50%, 6.51 N/mm) groups; the adjustable system was used for the dynamically adjusted group, in the 50% configuration (4.33 N/mm) for the first 7 days and the 100% configuration (11.41 N/mm) thereafter.

| **Tested ExFix system** | **Image** | **Loading Profile<br>Force – displacement curve** | **Stiffness (N/mm)** |
|---|---|---|---|
| Adjustable (100% side parts) | 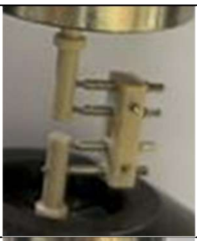 | 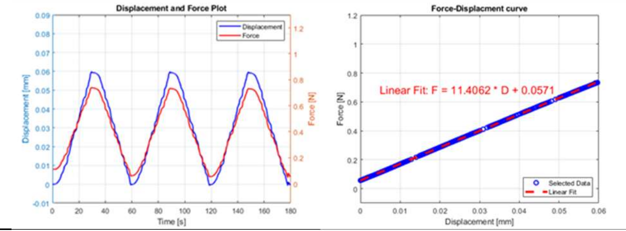 | **11.41** |
| Adjustable (50% side parts) | 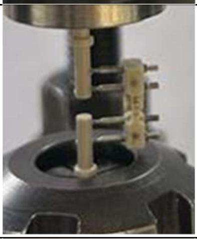 | 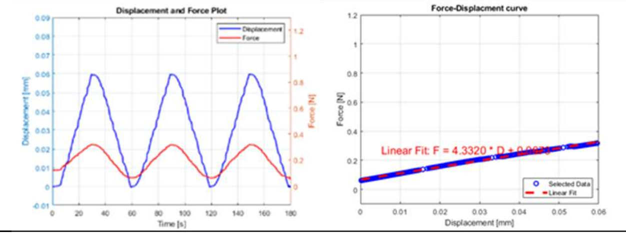 | **4.33** |
| Non-Adjustable (100%) | 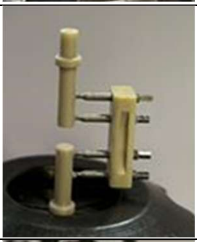 | 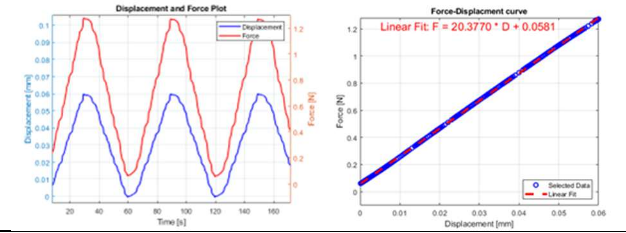 | **20.38** |
| Non-Adjustable (50%) | 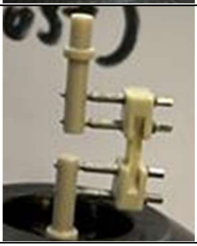 | 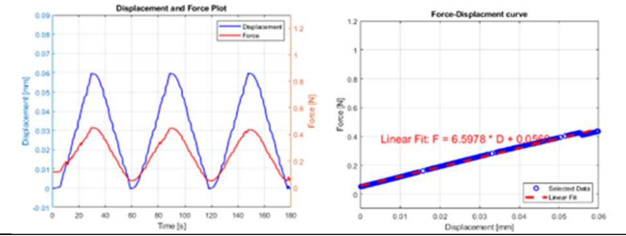 | **6.51** |

**Supplementary Table 2. Micro-computed tomography measurements at 21 days post-fracture.** Region of interest volume, total bone volume, and regenerated bone volume for each animal under rigid (n = 4), semirigid (n = 4) and dynamically adjusted (n = 3) fixation. Regenerated bone volume per ROI is expressed as a percentage in Figure 1c.

| Sample nr. | Fixation group | Roi ($mm^3$) | Bone_ROI ($mm^3$) | RegBone_ROI ($mm^3$) |
|---|---|---|---|---|

| | | | | |
|---|---|---|---|---|
| 1 | Rigid | 1.153174748 | 0.446034 | 0.409585 |
| 2 | Rigid | 1.434806405 | 0.649315 | 0.584728 |
| 3 | Rigid | 2.012402260 | 0.891709 | 0.752336 |
| 4 | Rigid | 1.333303484 | 0.721062 | 0.621249 |
| 5 | Semirigid | 1.155300840 | 0.336575 | 0.289328 |
| 6 | Semirigid | 2.890375990 | 0.86589 | 0.617209 |
| 7 | Semirigid | 2.661437454 | 0.922599 | 0.758126 |
| 8 | Semirigid | 1.836606183 | 0.648755 | 0.559813 |
| 9 | Adjustable | 1.302406540 | 0.807061 | 0.758446 |
| 10 | Adjustable | 1.143558054 | 0.646092 | 0.607555 |
| 11 | Adjustable | 1.249100319 | 0.756671 | 0.667968 |

**Supplementary Table 3. Micro-computed tomography measurements following Yoda1 treatment.** Region of interest volume, total bone volume and regenerated bone volume for each animal receiving Yoda1 or vehicle under rigid, semirigid and dynamically adjusted fixation. Treated animals were compared with vehicle controls of the same fixation condition by two-sided Welch's t-test. Rigid: 44.5 ± 2.9% versus 52.7 ± 3.5%, n = 3 and 3, p = 0.036 (Holm-adjusted p = 0.072), Hedges' g = 2.06. Semirigid: 24.5 ± 2.9% versus 24.7 ± 7.2%, n = 3 and 8, p = 0.94 (Holm-adjusted p = 0.94), Hedges' g = 0.03. Dynamically adjusted: 50.3 ± 3.2% versus 46.6 ± 20.9%, n = 2 and 5; not evaluated owing to the size of the vehicle group. Two comparisons were performed; p-values reported in the main text are uncorrected, as the three fixation conditions constituted independent cohorts.

| Sample nr. | Fixation group | Roi ($mm^3$) | Bone_ROI ($mm^3$) | RegBone_ROI ($mm^3$) |
|---|---|---|---|---|
| 1 | Rigid_Yoda1 | 1.764260365 | 1.022028 | 0.957312 |
| 2 | Rigid_Yoda1 | 1.221058261 | 0.747912 | 0.672527 |
| 3 | Rigid_Yoda1 | 0.743088510 | 0.376552 | 0.363530 |
| 4 | Rigid_CTRL | 1.076617783 | 0.563987 | 0.499315 |
| 5 | Rigid_CTRL | 2.597425630 | 1.506201 | 1.070222 |
| 6 | Rigid_CTRL | 1.555811593 | 0.916077 | 0.713684 |
| 7 | Semirigid_Yoda1 | 2.928727705 | 1.161865 | 0.867665 |
| 8 | Semirigid_Yoda1 | 2.180885353 | 0.560212 | 0.493913 |
| 9 | Semirigid_Yoda1 | 2.206191647 | 0.755551 | 0.60406 |
| 10 | Semirigid_Yoda1 | 2.693700897 | 0.941771 | 0.856108 |
| 11 | Semirigid_Yoda1 | 3.742864154 | 0.961663 | 0.759990 |
| 12 | Semirigid_Yoda1 | 1.446022528 | 0.229508 | 0.139821 |
| 13 | Semirigid_Yoda1 | 2.406118985 | 0.827121 | 0.722293 |
| 14 | Semirigid_Yoda1 | 2.861429212 | 0.980988 | 0.756903 |
| 15 | Semirigid_CTRL | 1.23659758 | 0.328473 | 0.279378 |
| 16 | Semirigid_CTRL | 2.086440421 | 0.691291 | 0.580728 |
| 17 | Semirigid_CTRL | 2.630949202 | 0.703945 | 0.606532 |

| | | | | |
|---|---|---|---|---|
| 18 | Adjustable_Yoda1 | 1.491195957 | 1.100965 | 0.998464 |
| 19 | Adjustable_Yoda1 | 1.365503528 | 0.736083 | 0.616946 |
| 20 | Adjustable_Yoda1 | 1.80795274 | 0.30708 | 0.221609 |
| 21 | Adjustable_Yoda1 | 1.858627443 | 1.015131 | 0.926517 |
| 22 | Adjustable_Yoda1 | 2.387390915 | 1.489031 | 1.403105 |
| 23 | Adjustable_CTRL | 2.015364282 | 1.080541 | 0.967189 |
| 24 | Adjustable_CTRL | 2.376915273 | 1.419542 | 1.248994 |

**Supplementary Table 4. Micro-computed tomography measurements following GsMTx4 treatment.** Region of interest volume, total bone volume and regenerated bone volume for each animal receiving GsMTx4 or vehicle under rigid, semirigid and dynamically adjusted fixation. Treated animals were compared with vehicle controls of the same fixation condition by two-sided Welch's t-test. Rigid: 43.7 ± 4.8% versus 18.1 ± 9.0%, n = 4 and 4, p = 0.005 (Holm-adjusted p = 0.010), Hedges' g = −3.11. Semirigid: 28.8 ± 4.6% versus 17.9 ± 10.2%, n = 4 and 4, p = 0.12 (Holm-adjusted p = 0.12), Hedges' g = −1.19. Dynamically adjusted: 57.0 ± 6.6% versus 14.9 ± 5.0%, n = 3 and 3, p = 0.001 (Holm-adjusted p = 0.004), Hedges' g = −5.76. Three comparisons were performed; p-values reported in the main text are uncorrected, as the three fixation conditions constituted independent cohorts.

| Sample nr. | Fixation group | Roi ($mm^3$) | Bone_ROI ($mm^3$) | RegBone_ROI ($mm^3$) |
|---|---|---|---|---|
| 1 | Rigid_GsMtx4 | 1.126219658 | 0.409121 | 0.354396 |
| 2 | Rigid_GsMtx4 | 1.919492495 | 0.389733 | 0.280314 |
| 3 | Rigid_GsMtx4 | 1.977498114 | 0.352276 | 0.254095 |
| 4 | Rigid_GsMtx4 | 1.494989299 | 0.276667 | 0.199402 |
| 5 | Rigid_CTRL | 0.903913705 | 0.416280 | 0.379055 |
| 6 | Rigid_CTRL | 1.179711919 | 0.599749 | 0.525067 |
| 7 | Rigid_CTRL | 1.921011817 | 0.871841 | 0.740962 |
| 8 | Rigid_CTRL | 1.427494743 | 0.794343 | 0.713087 |
| 9 | Semirigid_GsMtx4 | 2.981100072 | 0.333408 | 0.230148 |
| 10 | Semirigid_GsMtx4 | 1.249217996 | 0.427278 | 0.324538 |
| 11 | Semirigid_GsMtx4 | 1.677127451 | 0.603140 | 0.459864 |
| 12 | Semirigid_GsMtx4 | 2.180303860 | 0.298327 | 0.23022 |
| 13 | Semirigid_CTRL | 1.288512914 | 0.578257 | 0.451065 |
| 14 | Semirigid_CTRL | 2.187019073 | 0.842310 | 0.577377 |
| 15 | Semirigid_CTRL | 1.339761822 | 0.472325 | 0.392589 |
| 16 | Semirigid_CTRL | 0.976231438 | 0.312972 | 0.238210 |
| 17 | Adjustable_GsMtx4 | 1.395807907 | 0.329481 | 0.249784 |
| 18 | Adjustable_GsMtx4 | 0.839048863 | 0.119281 | 0.077257 |
| 19 | Adjustable_GsMtx4 | 1.052918909 | 0.302718 | 0.186100 |
| 20 | Adjustable_CTRL | 0.979005244 | 0.677238 | 0.630727 |
| 21 | Adjustable_CTRL | 1.058602078 | 0.735219 | 0.548071 |
| 22 | Adjustable_CTRL | 1.592595418 | 0.974997 | 0.871689 |

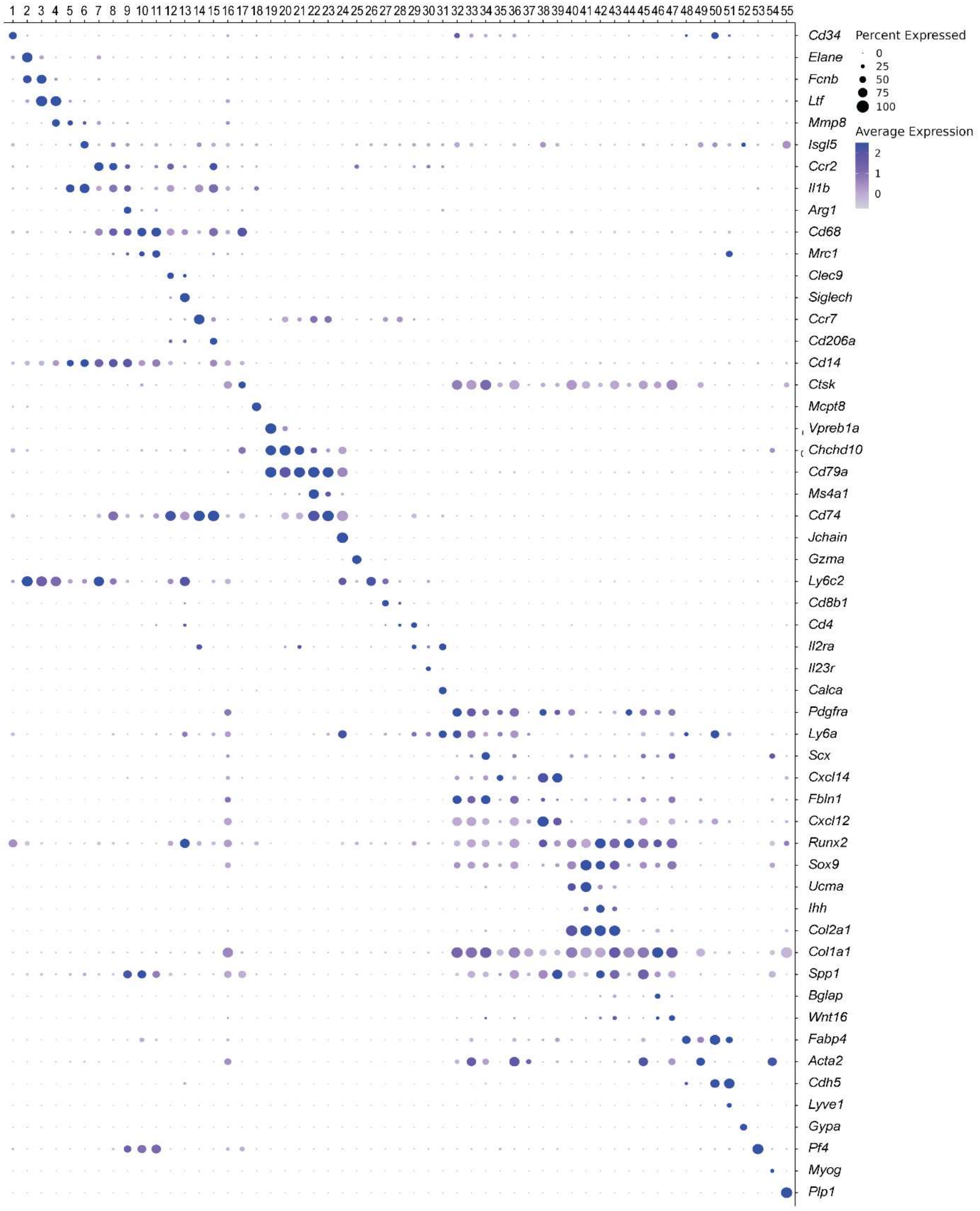


**Supplementary Figure 1. Marker gene expression across annotated cell populations.** Dot plot showing expression of selected marker genes used to annotate the 55 populations identified in the integrated single-cell dataset. Dot size indicates the fraction of cells expressing each gene; color indicates scaled mean expression.

1: Granulocyte Monocyte Precursor, 2: G1 Neutrophil, 3: G2 Neutrophil, 4: G3 Neutrophil, 5: G4 Neutrophil, 6: G5 Neutrophil, 7: Monocyte, 8: *Il1b*+ Macrophage, 9: *Arg1*+ Macrophage, 10: *Cd68*+ Macrophage, 11: *Mrc1*+ Macrophage, 12: *Clec9a*+ DC, 13: pDC, 14: Migratory DC, 15: moDC, 16: Fibrocyte, 17: Osteoclast, 18: Mast Cell, 19: Late Pro-B Cell, 20: Pre-B Cell, 21: Immature B Cell, 22: Naive B Cell, 23: Activated B Cell, 24: Plasmacell, 25: Natural Killer Cell, 26: γδ T Cell, 27: *Cd8*+ T Cell, 28: *Cd4*+ T Cell, 29: Treg Cell, 30: Th17 Cell, 31: ILC2, 32: pSSPC, 33: bmSSPC, 34: TSPC, 35: Fibroblast Progenitor, 36: Proliferating Fibroblast, 37: Fibroblast, 38: Adipo-CAR, 39: Osteo-CAR, 40: Chondroprogenitor, 41: Resting Chondrocyte, 42: Hypertrophic Chondrocyte, 43: Osteoblast-Committed Chondrocyte, 44: Low-Activity Osteoprogenitor, 45: Pre-Osteoblast, 46: Osteoblast, 47: Periosteum, 48: Adipocyte, 49: Perivascular Cell, 50: Vascular Endothelial Cell, 51: Lymphatic Endothelial Cell, 52: Erythroid Cell, 53: Megakaryocyte, 54: Myogenic Cell, 55: Neuron

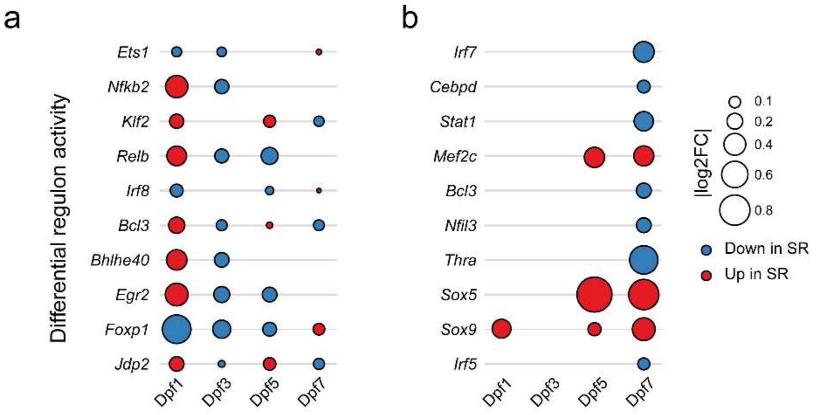


**Supplementary Figure 2. Differential regulon activity across all timepoints.** Gene regulatory network analysis comparing semirigid and rigid fixation in immune populations **a**, and mesenchymal and endothelial populations **b**, on days 1, 3, 5 and 7 post-fracture. Dot size indicates the absolute log2 fold change in regulon activity; color indicates the condition in which activity was higher.

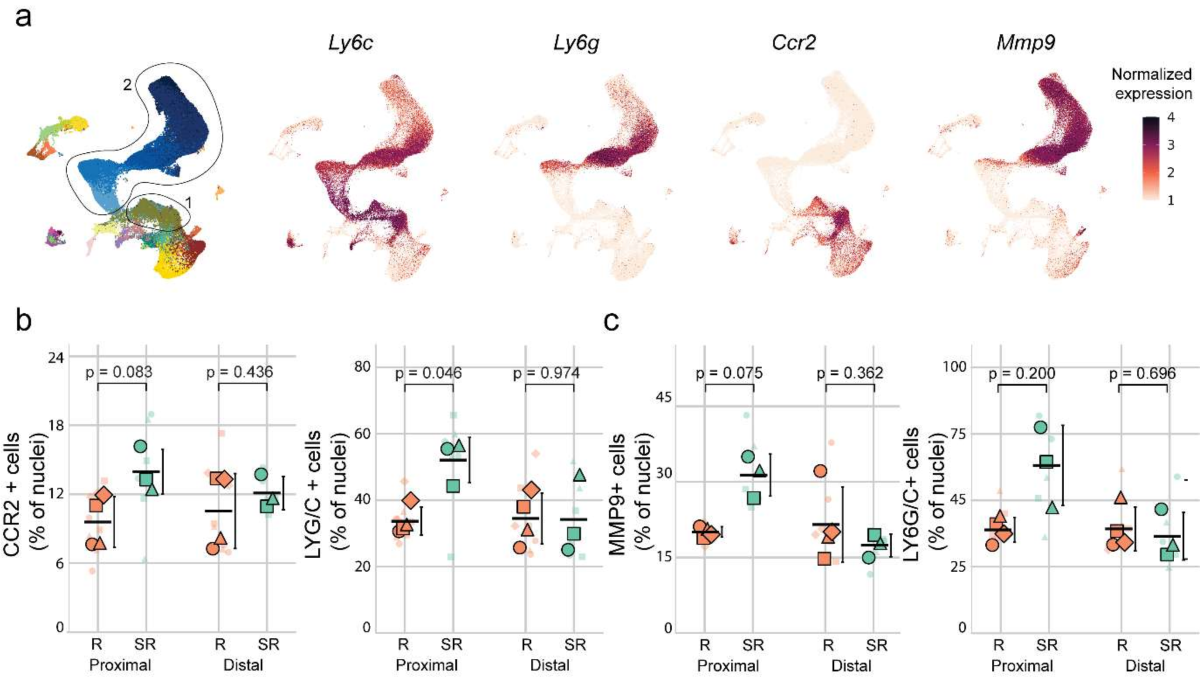


**Supplementary Figure 3. Marker expression and immunofluorescence quantification of monocytes and neutrophils on day 1. a**, Left, UMAP of immune populations across all timepoints, colored by cell type as in Figure 2a, with outlined regions corresponding to monocyte (1) and neutrophil (2) populations. Right, feature plots showing expression of *Ly6c*, *Ly6g*, *Ccr2* and *Mmp9* **b**, CCR2+ (left) and LY6G/C+ (right) cells as percentage of nuclei in proximal and distal regions of the fracture callus in the LY6G/C/CCR2 double staining (Figure 2g). **c**, MMP9+ (left) and LY6G/C+ (right) cells as percentage of nuclei in the LY6G/C/MMP9 double staining (Figure 2h). In b and c, small symbols show individual regions of interest (two sections per animal) and large symbols animal means, with symbol shape identifying the animal; horizontal lines show group means with SD. Rigid (R) and semirigid (SR) fixation were compared within each region by Welch's t-test with Holm correction for two comparisons. n = 4 rigid and 3 semirigid animals.

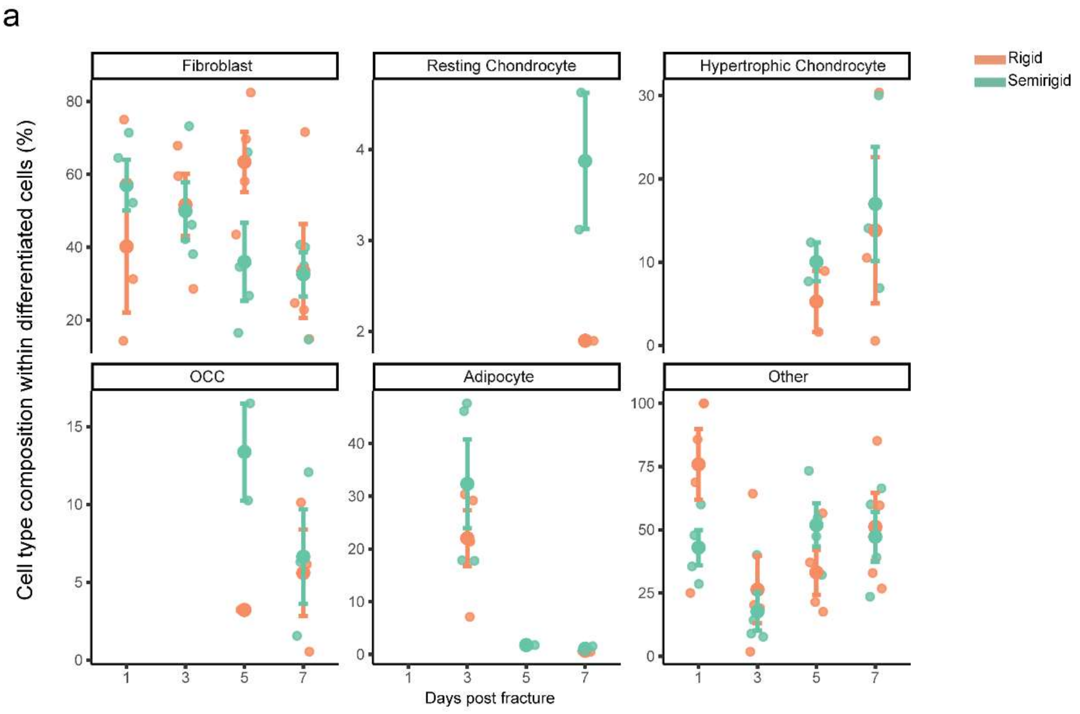


**Supplementary Figure 4. Cell type composition within differentiated populations across the first week.** Proportions of differentiated cell types under rigid and semirigid fixation on days 1, 3, 5 and 7 post-fracture. Given the limited abundance of several populations, these are descriptive distributions rather than statistically supported compositional shifts.

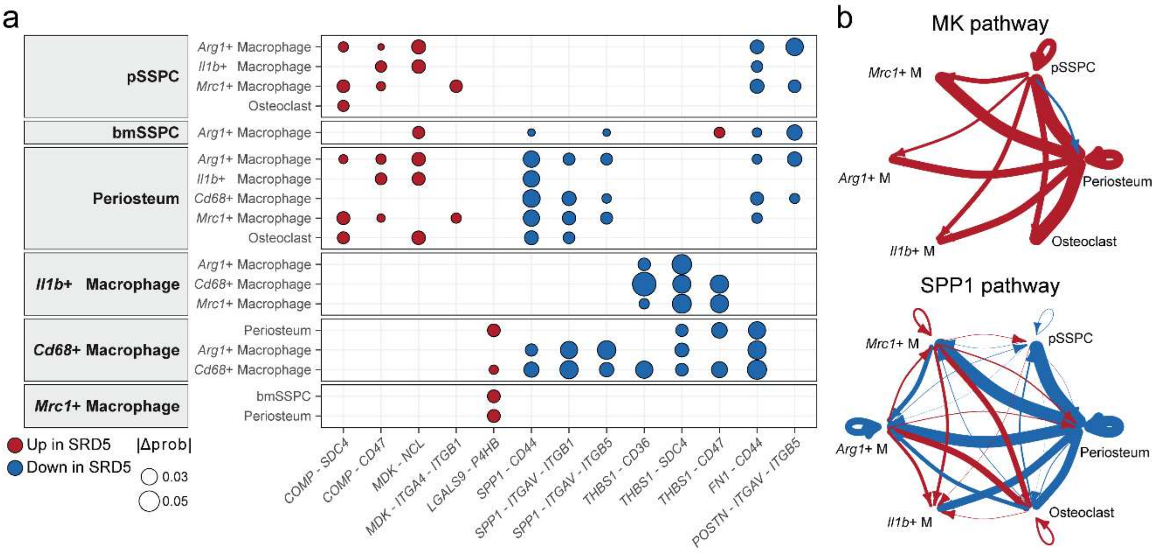


**Supplementary Figure 5. Differential cell–cell communication on day 5. a**, Predicted ligand–receptor interactions between skeletal progenitor, periosteal and macrophage populations under semirigid compared with rigid fixation at 5 days post-fracture. Senders are indicated on the left, receivers on the vertical axis. Dot color indicates interactions increased under semirigid (red) or rigid (blue) fixation; dot size indicates the magnitude of the change in communication probability (|Δ probability|). **b**, Differential communication networks for the MK (top) and SPP1 (bottom) pathways between the same populations. Arrow direction indicates sender to receiver; width is proportional to the difference in communication probability; color indicates direction as in **a**. Populations shown are those present at ≥25 cells in both conditions with ≤3-fold difference in abundance. Total network information flow was closely matched between conditions at this timepoint. n = 4 biological replicates per condition. SRD5: semirigid day 5, RD5: rigid day 5.

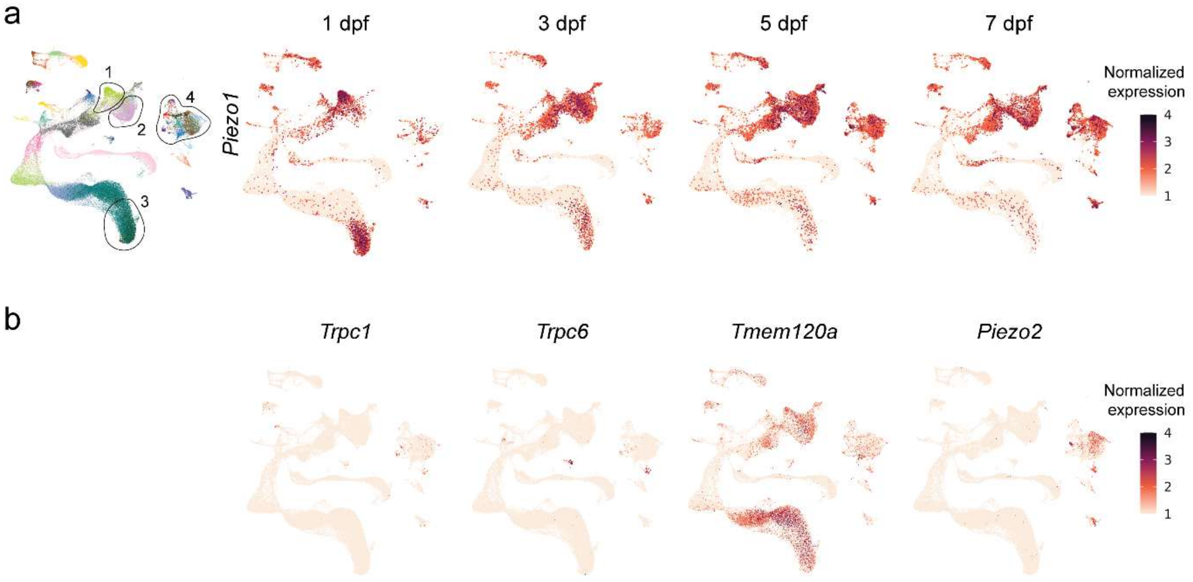


**Supplementary Figure 6. Expression of mechanotransduction-associated genes across the fracture callus. a**, Left, reference UMAP with the four regions referred to in the main text outlined: 1: *Arg1*+ Macrophages, 2: *Mrc1*+ Macrophages, 3: G5 Neutrophils, 4: Mesenchymal Cells. Right, *Piezo1* expression at 1, 3, 5 and 7 days post-fracture. **b**, Expression of documented alternative targets of GsMTx4, *Trpc1*, *Trpc6*, *Tmem120a* and *Piezo2*, projected onto the same UMAP. n = 4 to 5 biological replicates per condition and timepoint; rigid and semirigid samples pooled. Dpf: days post-fracture.

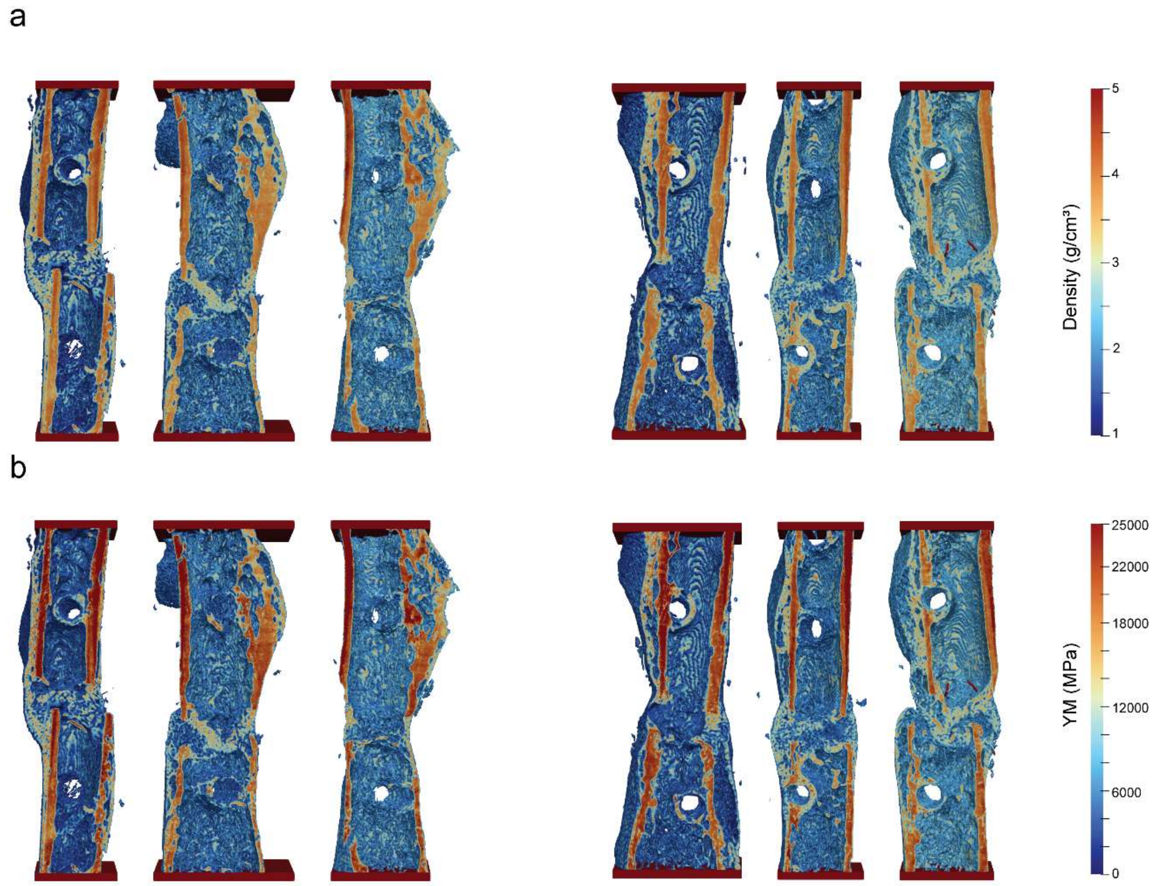


**Supplementary Figure 7. Finite element models of healed calluses following Yoda1 treatment.** **a**, Tissue mineral density distributions derived from micro-computed tomography for vehicle-treated (left) and Yoda1-treated (right) specimens under rigid fixation at 21 days post-fracture. **b**, Corresponding Young's modulus distributions for the same specimens, derived from the density values in panel a. Color scales indicate density (g/cm$^3$) and Young's modulus (MPa). Red bands indicate the loading plates applied in the simulated torsional loading. n = 3 specimens per group.

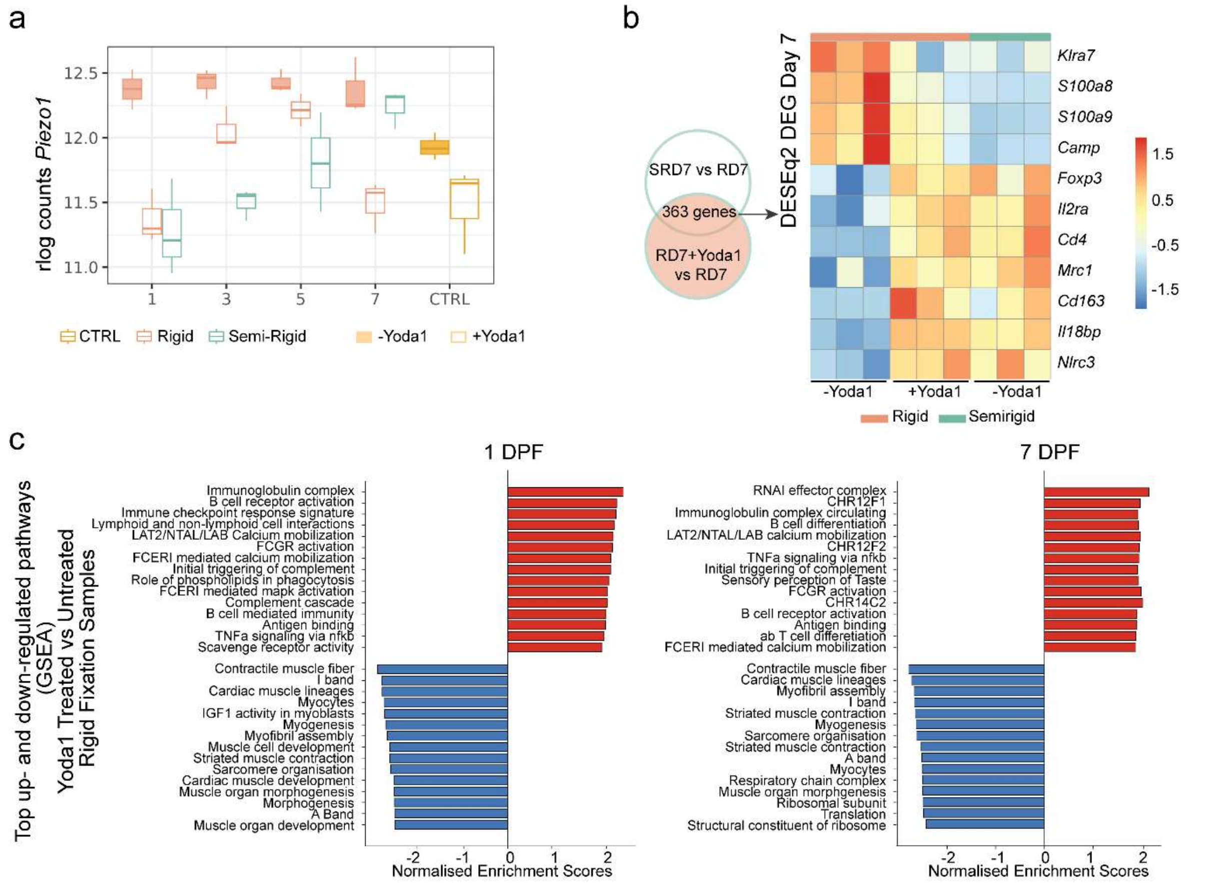


**Supplementary Figure 8. Transcriptional response to PIEZO1 activation. a**, *Piezo1* expression in fracture calluses under rigid and semirigid fixation on days 1, 3, 5 and 7 post-fracture, and in Yoda1-treated and untreated rigid calluses (rlog-normalized counts). CTRL, unfractured bone. **b,** Left, overlap between genes differentially expressed between semirigid and rigid fixation (SRD7 versus RD7) and between Yoda1-treated and untreated rigid fixation (RD7+Yoda1 versus RD7) at 7 days post-fracture. Right, expression of selected shared genes across all groups; color indicates row-scaled z-score of DESeq2-normalized counts, with each row scaled independently. **c**, Gene set enrichment analysis of Yoda1-treated compared with untreated rigid fixation samples at 1 and 7 days post-fracture, showing the most strongly up- and down-regulated gene sets by normalized enrichment score. n = 3 biological replicates per condition. Yoda1-treated samples were sequenced on a different platform from the untreated samples; this comparison is therefore descriptive.
